# Genomic and immunogenic changes over the history of the Viral Hemorrhagic Septicemia (VHS-IVb) fish virus (=*Piscine novirhabdovirus*) in the Laurentian Great Lakes

**DOI:** 10.1101/2020.04.27.063842

**Authors:** Megan D. Niner, Carol A. Stepien, Bartolomeo Gorgoglione, Douglas W. Leaman

**Author notes:** All Correspondence; m. Joint First Authors contributed equally to this work (Megan D. Niner, Carol A. Stepien).

## Abstract

Viral Hemorrhagic Septicemia Virus (VHSV) (=*Piscine novirhabdovirus*) appeared in the Laurentian Great Lakes in 2005, constituting a unique and highly virulent genogroup (IVb), which killed >32 fish species in large 2005 and 2006. Periods of apparent dormancy punctuated smaller outbreaks in 2007, 2008, and 2017. We conducted the first whole genome analysis of IVb, evaluating its evolutionary changes using 46 isolates, in reference to immunogenicity in cell culture, and the genomes of other VHS genogroups (I–IVa) and other Novirhabdoviruses. IVb isolates had 253 genomic nucleotide substitutions (2.3% of the total 11,158 nucleotides), with 85 (16.6%) being non-synonymous. The greatest number of substitutions occurred in the non-coding region (NCDS; 4.3%) followed by the *Nv-* (3.8%), and *M-* (2.8%) genes. The *M*-gene possessed the greatest proportions of amino acid changes (52.9%), followed by the *Nv-* (50.0%), *G-* (48.6%), *N-* (35.7%) and *L-* (23.1%) genes. Among VHS genogroups, IVa from the northeastern Pacific exhibited the fastest substitution rate (2.01×10-3), followed by Ivb (6.64×10^−^5), and I/III from Europe (4.09×10^−^5). A 2016 gizzard shad isolate from Lake Erie was the most divergent IVb isolate (38 NT, 15.0%, 15 AA), yet exhibited reduced virulence with *in vitro* immunogenicity analyses, as did other 2016 isolates, in comparison to the first IVb isolate (2003). The 2016 isolates exhibited lower impact on innate antiviral responses, suggesting phenotypic effects. Results suggest continued sequence change and lower virulence over the history of IVb, which may facilitate its long-term persistence in fish host populations.

## Introduction

Viral Hemorrhagic Septicemia Virus (VHSV), a.k.a. *Piscine novirhabdovirus*, causes one of the world’s most serious viral finfish diseases, infecting >140 aquaculture and wild fish species across the northern hemisphere [1,2]. Like many other RNA viruses, VHSV has a short genome, lack of proofreading, and rapid generation time [3,4,5]. These factors may lead to its rapid evolution, facilitating adaptation to new hosts and environments [6,7,8].

VHSV suddenly appeared in North America’s Laurentian Great Lakes in 2005, causing large fish kills of >32 species in 2005-6 [9,10,11]. It was designated as the geographically and genetically distinct subgenogroup (=substrain) IVb. Prior to this, VHS was known as three genogroups (=strains) in Europe (I-III; dating to the 1930s), and genogroup IVa in the northeastern Pacific (since the 1980s) and northwestern Pacific (1990s) (summarized by [12]).

Since its emergence, VHSV IVb outbreaks have become less prevalent [13,14,15], interspersed by small, geographically restricted occurrences in 2007, 2009, 2011, and 2017 [13, 16](M. Faisal, personal communication 2017; R. Getchell personal communication 2017). Yet, the virus has continued to diversify, according to sequences of selected gene regions [14]. Its diversification appears to have followed a “quasispecies” evolutionary pattern of multi-directional, “cloud-like” diversification of closely related variants [7,8,12]. In the present study, we sequenced the entire genome of available isolates to evaluate its evolutionary patterns over time and across its geographic range. We relate these to laboratory culture virulence studies. Knowledge of evolutionary changes across VHSV-IVb’s genome and the functional roles of genes will aid understanding of the co-evolutionary “arms race” between this pathogen and its hosts.

### VHSV genome and evolutionary history

VHSV has a single-stranded, negative sense RNA genome of 11,158 nucleotides (NTs), and does not recombine [17, 18]. It contains five structural genes: nucleoprotein (*N*), phosphoprotein (*P*), matrix (*M*), glycoprotein (*G*), and large protein (*L*) 5’*N*–*P*–*M*–*G*–*Nv*–*L*’3. VHSV belongs to the genus *Novirhabdovirus*, whose species also include *Salmonid novirhabdovirus* (IHNV), *Hirame novirhabdovirus* (HIRRV) and *Snakehead novirhabdovirus* (SHRV). All of these infect fishes, and are distinguished by their unique non-virion (*Nv*) gene. *Nv* encodes a distinctive non-structural protein [2, 18, 19], whose evolutionary history was unstudied prior to our investigation.

The functionality of the unique *Nv*-gene has been of scientific conjecture, which varies among the four species. Although non-essential for replication, *Nv* is required for pathogenicity in VHSV [19] and IHNV [20], but not in SHRV [21, 22]. Interchanging *Nv* between VHSV and IHNV reduced pathogenicity in both recombinants [23]. *Nv* appears to function in suppressing host innate immune response (IIR) [23, 24, 25] and preventing apoptosis [19], but its exact mechanisms are unknown. Single sequence changes in *Nv* can impact the capacity to suppress host cell responses [26, 27]. Although *Nv* characterizes Novirhabdoviruses, its sequence is not conserved among species [18], and our literature search found no prior investigations of its potential evolutionary origin; which is tested here.

VHSV comprises four geographically and genetically distinct lineages: I–III in Europe and IV in North America and Asia (Figures 1 and 2) [12, 28, 29]. VHSV-I is the oldest-discerned, possessing the largest known geographic range, the most named phylogenetic subgroups, and greatest number of described host species [12, 15, 18]. VHSV-I infections have caused significant economic losses to European aquaculture [30, 31].

**Figure 1.**
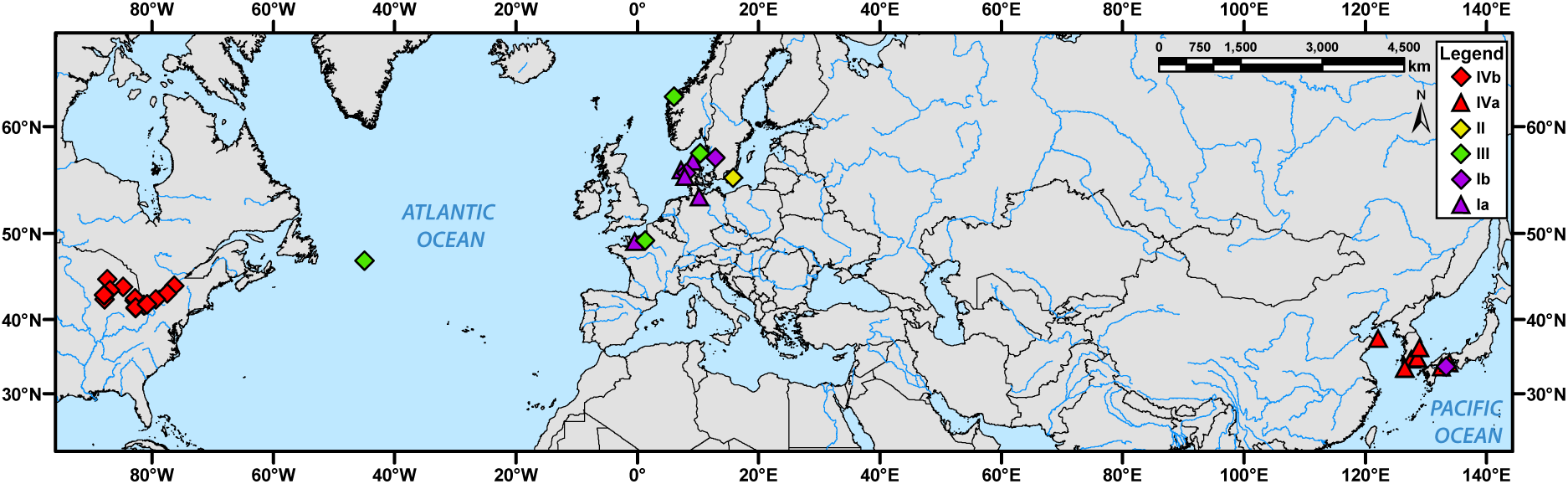
Map of VHSV (I–IV) full genome isolates included in this study, colored by Genogroup and with shapes denoting substrains.

**Figure 2.**
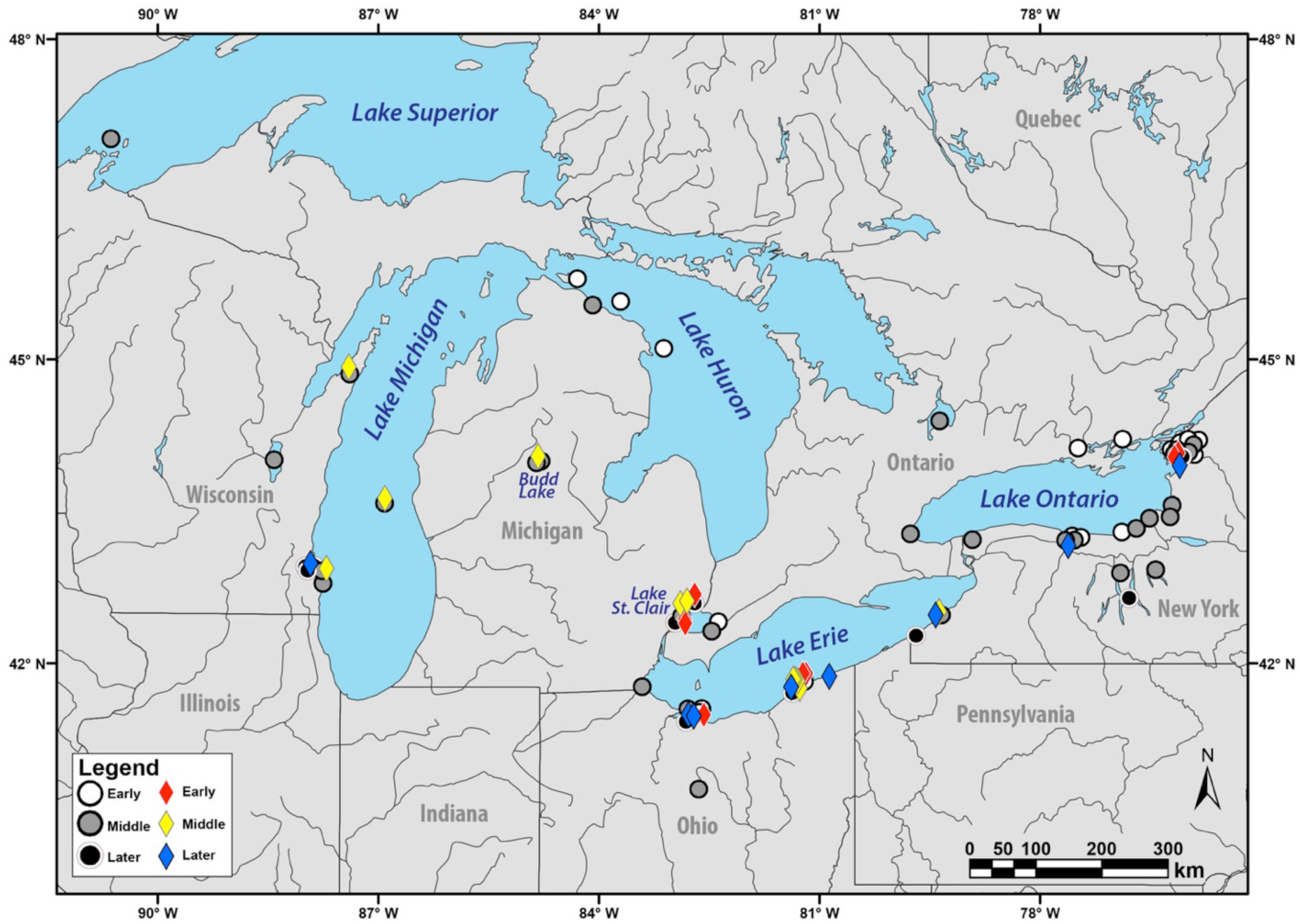
Map of VHSV-IVb occurrences in the Great Lakes. Shapes colored for time periods (Early: 2003–06, Middle: 2007–10, Late: 2011–17). Diamonds are isolates with sequenced genomes. Large circles depict VHSV-IVb that lack whole genome data.

Phylogenetic analyses found that I, II, and III shared common ancestry, separate from genogroup IV [12, 14, 15]. VHSV-IV contains three described genogroups, with IVa initially described from the northeastern Pacific coastal region in the 1980s [32, 33, 34]. In the late 1990s, IVa also appeared along the Asian Pacific coast, off Korea and Japan [35]. VHSV-IVc occurs in northwestern Atlantic marine waters, dating to its discovery in 2000 [36]. Phylogenetic analyses have placed IVc as the sister group to IVb, which together (IVb–c) then comprises the sister group to IVa [12, 14,15]. An unclassified potential fourth genogroup, appearing derived from IVa, was isolated from wild lumpfish (*Cyclopterus lumpus*) in Breidafjördur Bay, Iceland [37].

### VHSV infection and host immune responses

VHSV infection is characterized by petechial hemorrhaging diffuse to external and internal organs, with surviving fishes shedding virus up to three months post-infection [38]. Pathology studies largely have used the original VHSV-IVb 2003 isolate (C03MU) [39, 40, 41] or experimentally mutated versions [19, 42, 43], leaving uncertainty about the traits of new variants. Just two investigations have compared the original isolate to adaptive traits of other naturally occurring IVb isolates; Imanse et al. [44] found reduced virulence and lower mortality in a 2010 Lake Ontario isolate, and Getchell et al. [45] discerned lower viral titers with no mortality differences in three other round goby isolates.

After gaining entry to the host cell, the virus must successfully inhibit its innate immune defense in order to replicate. Cellular recognition of viral byproducts triggers several host signaling cascades, especially the interferon (IFN) pathway [46]. An array of Type I IFNs are highly induced by VHSV infection, triggering pathogenesis [47]. IFNs bind to specific surface receptors of effector cells, initiating transcription of IFN-stimulated genes to initiate antiviral response [48]. Thus, comparatively measuring IFN production provides insights on how effectively the virus slows down the host response, serving as a proxy of potential host-pathogen evolution.

### The Red Queen evolutionary “arms race”

Host-pathogen relationships often are termed an evolutionary “arms-race” described by Van Valen’s [49, 50] “Red Queen” hypothesis, in which selection pressures from pathogens lead either to death or adaptation in host populations, and *vice-versa*. Since RNA viruses mutate just below the maximum limit for functionality, they readily can adapt to host defense modulations [51]. VHSV-IVb has been mutating over time [11, 14, 15], however, little is known about how its more recent isolates behave during infection. The rising importance of the aquaculture industry has raised concerns about fish pathogens from economic and sustainability perspectives [52, 53].

Biotechnology developments, including high-throughput sequencing (HTS), now allow analyses of entire viral genomes to investigate evolutionary patterns in pathogen lineages [54]. Our objective here is to understand evolutionary relationships and trajectories across the VHSV-IVb genome, and evaluate spatial and temporal patterns in relation to host-pathogen coevolution. Only five VHSV-IVb genomes were previously fully sequenced [26, 45, 55], and just one other publication evaluated similarities and differences based on those few isolates [45]. Ours is the first study to compare phylogenetic trends among entire VHSV-IVb genomes, analyzing isolates across the Great Lakes throughout their history and geographic range, and evolutionary patterns among genes. We compare the evolutionary patterns of IVb variants to other VHSV genogroups and Novirhabdoviruses. We additionally examine pathology changes of 2016 VHSV-IVb isolates to the initial 2003 isolate, in relation to IFN induction and virulence.

## Materials and methods

### Sampling and nomenclature

Samples from 2015–17 were obtained by from Niner and Stepien [see 56]. Historical VHSV-IVb isolates were provided by G. Kurath (USGS, Seattle, WA) as frozen infected media from Bluegill Fry (BF-2) cell culture. Additional complete VHSV (I–IVa; Table 1) and Novirhabdovirus (Table 2) genomes were downloaded from GenBank and aligned using MEGA X. VHSV-IVb isolates are named here with unique identifiers, by first letter of lake name, the last two digits of its isolation year, followed by the first two letters of the host species’ common name. Example: the original isolate, collected from a Lake St. Clair muskellunge in 2003 here is C03MU. If more than one isolate shared the above identifiers, an additional lower-case letter was added (Table 1). We defined a haplotype as “a unique gene sequence differing by one or more nucleotide substitutions” from C03MU.

**Table 1.**
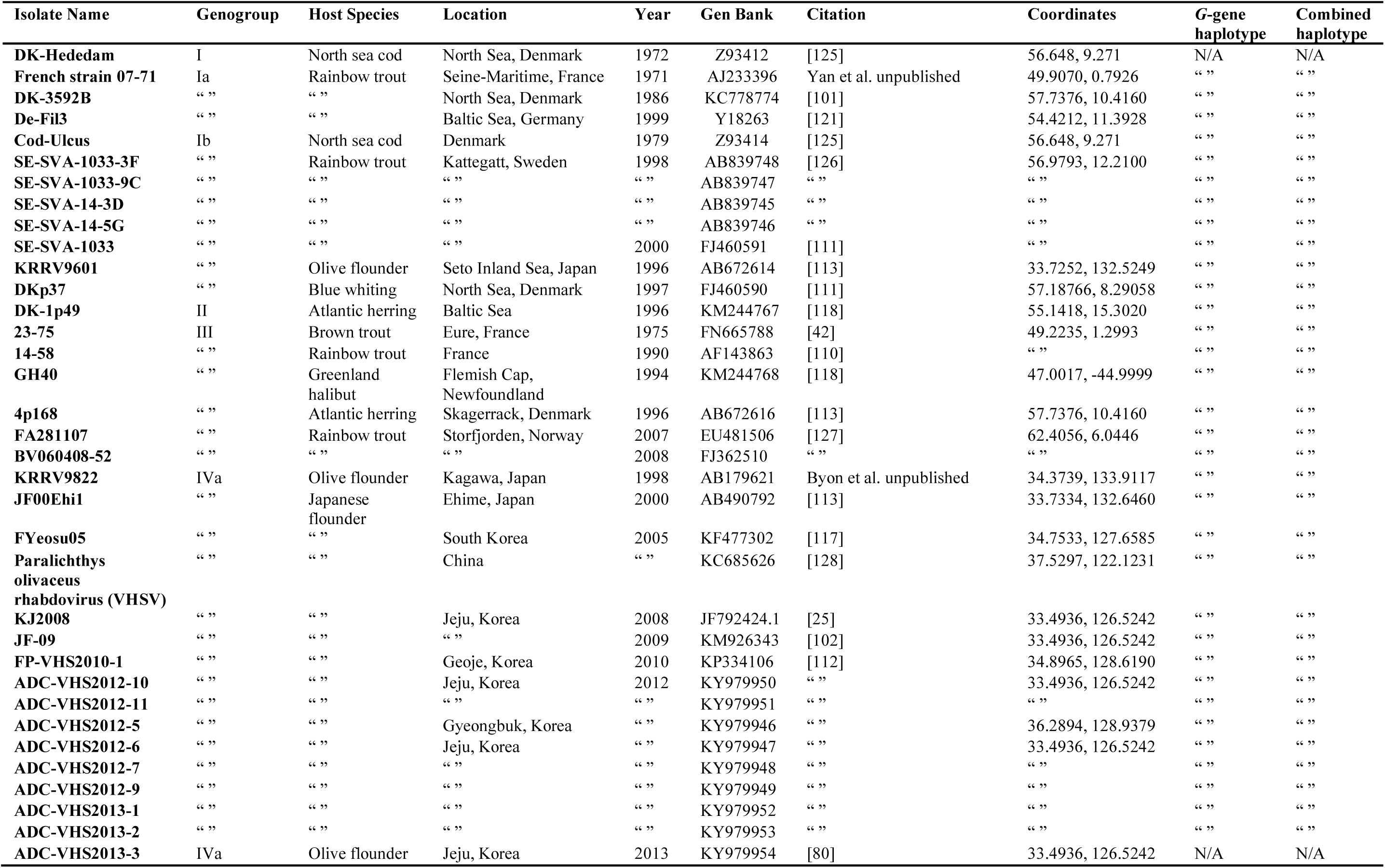
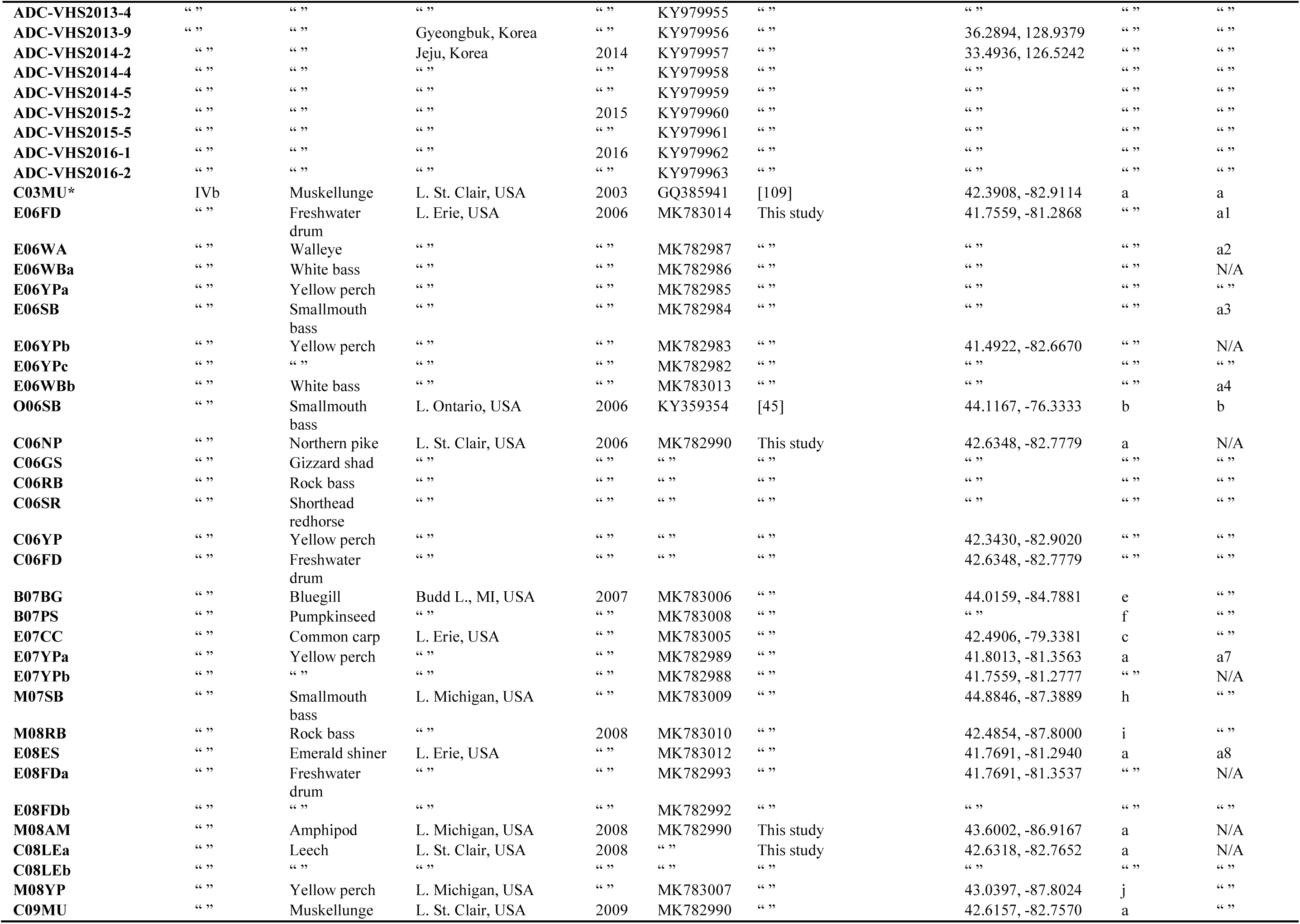
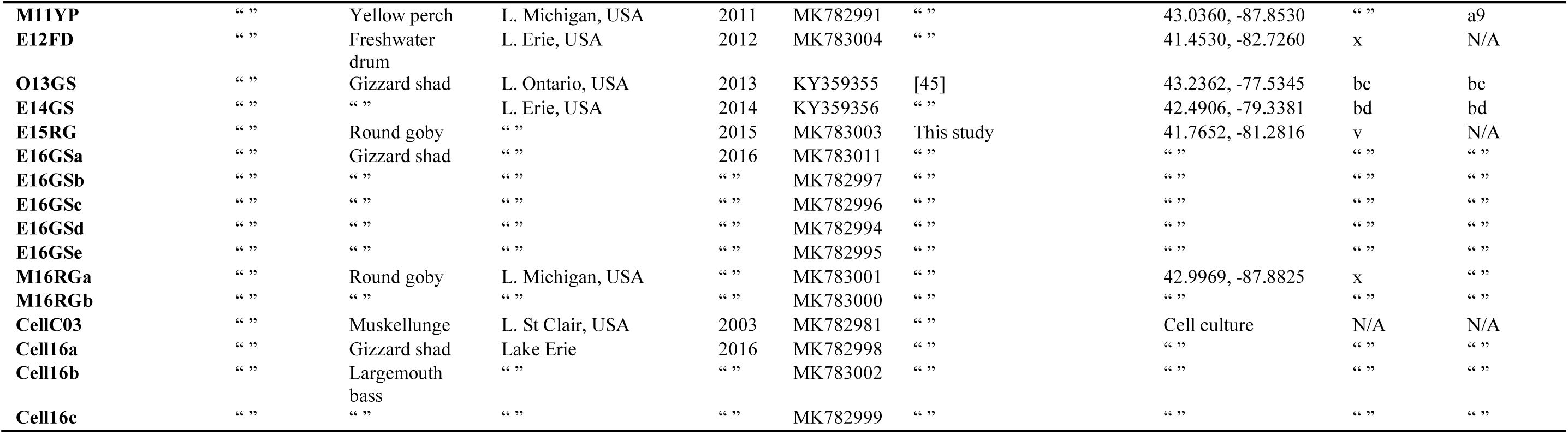
VHSV isolates for full genome analysis. IVb isolates analyzed and their corresponding *G*-gene and combined gene haplotypes.

**Table 2.** Additional Rhabdovirus sequences used in phylogenetic trees.

| Isolate Name | Host | Location | Year | GenBank | Sequence Citation |
| --- | --- | --- | --- | --- | --- |
| **Infectious Hematopoietic Necrosis Virus (IHNV)** |  |  |  |  |  |
| **X89213** | Rainbow trout | Oregon, USA | 1969 | X89213 | [121] |
| **WRAC** | Chinook salmon | Idaho, USA | 1994 | L40883 | [119] |
| “ ” | “ ” | “ ” | “ ” | NC_001652 | “ ” |
| **220-90** | Rainbow trout | Idaho, USA | 1990 | GQ413939 | [109] |
| **HLJ-09** | “ ” | China | 2009 | JX649101 | [122] |
| **Ch20101008** | Brook trout | “ ” | 2010 | KJ421216 | [114] |
| **BjLL** | Rainbow trout | “ ” | 2012 | MF509592 | [124] |
| **Snakehead Rhabdovirus (SHRV)** |  |  |  |  |  |
| **NC_000903** | Snakehead murrel | Thailand | 1988 | NC_000903 | [115] |
| **AF147498** | “ ” | “ ” | “ ” | AF147498 | [21] |
| **Hirame rhabdovirus (HIRRV)** |  |  |  |  |  |
| **CA 9703** | Japanese flounder | Japan | 1984 | NC_005093 | [116] |
| “ ” | “ ” | “ ” | “ ” | AF104985 | “ ” |
| **80113** | Stone flounder | China | 2008 | FJ376982 | [123] |

### Virus isolation

RNA was extracted using a previously optimized Trizol protocol [57], re-suspended in RNase-free water, and quantified with a NanoDrop 2000 spectrophotometer (Thermo Fisher Scientific). Frozen media was thawed on ice, and 30 µL from each sample was diluted 1:5 with serum-free MEM (ThermoFisher Scientific), and added to individual wells of a 12 well plate with confluent BF-2 monolayers. Cells were incubated 1 hr with the infected media at 20°C, then replaced with complete MEM 10% (v/v) cosmic calf serum (GE Healthcare) and 1% penicillin/streptomycin antibiotics (Invitrogen). Infected cells were incubated at 20°C for ≤one week and sampled when ≥80% CPE was achieved. Media was collected and added to a 1.5ml tube with 250 µL of versene (Fisher Scientific) for 10 min prior to 4 min 4000 rpm centrifugation at 4°C. Supernatant was discarded, and the versene/cell mixture spun again as before. Remaining supernatant was discarded again, and 250 µL Trizol® (ThermoFisher Scientific) was added to the remaining pellet.

### Genome sequencing

cDNA was synthesized from total RNA extracted from tissue samples using SuperScript IV (Invitrogen), following manufacturer’s instructions. Genomic cDNA was amplified in four segments using primers from Schönherz [58], substituting VHSV_Frag1I_nt18_+s with a more specific primer (5’GAGAGCTGCAGCACTTCACCG C3’), and 1µl cDNA in 25µL reactions with One Taq DNA polymerase (New England Biolabs). Amplicons were examined under UV light on 1% agarose gels stained with ethidium bromide. Target PCR products were gel excised and purified using QIAquick Gel Extraction kits (Qiagen).

Additional PCRs amplified the front 700 nucleotides (NTs) and end 400 NTs of the genome, with 45 s extension time. The front segment utilized VHSV_Frag1I_nt18_+s (5’GAGTTATGTTACARGGGACAGG3’) [58] and anti-sense 5’TGACCGAGATGGCAGATC3’, and end primers were designed based on the VHSV-IVb genome (GenBank: GQ385941) (End sense: 5’CCCAGATGCTATCACCGAGAA3’, End anti-sense 5’ACAAAGAATCCGAGGCAGGAG3’). Cleaned products were Sanger sequenced at Cornell DNA Services (Ithaca, NY) and sequences aligned and analyzed by us using MEGA X [59].

Genomic sequencing was outsourced to Ohio State University’s Molecular and Cellular Imaging Center (Wooster, OH). Sequences were uploaded by us to the Galaxy web platform and analyzed with usegalaxy.org programs [60]. Segments were aligned to the reference VHSV-IVb genome (C03MU, GenBank: GQ385941) using MAP WITH BWA-MEM [61]. For each of the 46 isolates, consensus sequences were generated followed by manual checking of each single nucleotide polymorphism (SNP) and coverage read using Integrative Genomics Viewer (IGV) [62, 63]. Consensus sequences, front, and end segments were concatenated, aligned, and trimmed in MEGA X [59].

### Genetic analyses

The Basic Local Alignment Search Tool (BLAST) was used to detect similar sequences to *Nv*, excluding the *Novirhabdovirus* genus. JMODELTEST v3.7 [64] employed the Akaike Information Criterion (AIC) to determine best-fit evolutionary models [65]. Phylogenetic analyses evaluated the most parsimonious evolutionary relationships (per [12, 14]) with Maximum Likelihood (ML) (PHYML v3.0) [66] and Bayesian (MRBAYES v3.1) [67] analyses. For the latter, Metropolis-coupled Markov Chain Monte Carlo (MCMCMC) analyses were run for five million generations, sampling every 100 to obtain posterior probability values. Burn-in was determined by plotting log likelihood values to identify stationarity, discarding the first 25%. Branch support for ML was calculated from non-parametric bootstrapping replications, which were limited to 500 for the respective *Novirhabdovirus*, 1450 for IVb, 2000 for *Nv*, and 500 for the *Nv* amino acid (AA) trees.

Haplotype networks were analyzed in POPART (www.popart.otago.ac.nz) using TCS [68]. We discerned whether samples significantly diverged over time or space using pairwise *θ*_ST_ (*F*_ST_ analogue) [69] comparisons in ARLEQUIN [70], reporting values prior and after sequential Bonferroni corrections [71].

### Evaluating evolution and selection

Comparative divergence times were estimated using BEAST v1.10.4 [72], with JMODELTEST output, a relaxed molecular clock with lognormal distribution, sampled every 50,000 of 500,000,000 generations. Outputs were assessed with TRACER v1.5 (in BEAST) to ensure stationarity. Collection dates were used as calibration points and tree branches set following PHYML output. Numbers of nucleotide substitutions per site per year (*k* = substitutions site–1 yr–1) were determined from pairwise (p) distances. Nucleotide (NT) and amino acid (AA) substitutions were evaluated for all IVb isolates and compared to C03MU. IVa isolates were compared to KRRV9601 (GenBank: AB179621) and I/III was compared to the European Ia isolate Hededam (GenBank: Z93412).

Two codon-based methods examined the possibility of selection pressures for each gene. Fast, unconstrained Bayesian approximation (FUBAR) [73] identified positive or purifying selection; pressures selecting for beneficial traits. However, FUBAR’s assumption of constant selection may not accurately represent IVb, since selection pressures may differ across hosts and environmental factors. To remedy this, we used MEME (mixed effects model of evolution) [73], which can detect positive selection, under strong purifying selection or the removal of detrimental variants. FUBAR and MEME were run with HyPHY on DataMonkey (www.datamonkey.org), with significance evaluated using posterior probability >0.95 for FUBAR and *p*<0.05 for MEME.

Amino acid sequences were submitted to the Phyre2 web portal (https://www.sbg.bio.ic.ac.uk/phyre2/) for *G*-gene protein modeling, prediction, and analysis [74]. For open reading frame (ORF) analysis, full length cDNA nucleotide sequences per isolate were submitted to the NCBI ORF Finder (https://www.ncbi.nlm.nih.gov/orffinder/) to search for alternate products >300nt in length. Sequences returned coding in reverse were discounted since VHSV is single-stranded.

### Cell lines and cell culture experiments

*Epithelioma papulosum cyprinid* (EPC) (ATCC: CRL-2872) and BF-2 (ATCC: CCL-91) cells were cultured as above. Three viable independent viral stocks of 2016 isolates, E16GSa (Cell16a) and E16LB (Cell16b and Cell16c) were derived from pooled organs (kidney, liver, and spleen) of sampled fish. Cell culture amplified stocks are denoted as “Cell”. Due to low concentration, the original E16LB sample was just partially sequenced, matching E16GSa within those regions [75]. The cell culture amplified reference control (CellC03) was derived from C03MU. VHSV-IVb isolates were amplified for subsequent purification by infecting confluent monolayers of BF-2 cells in 15 cm tissue culture dishes with 1:1000 (v/v) dilution of un-purified virus stock in serum-free EMEM. Viral adsorption occurred for 1 h, then medium was replaced with complete EMEM. Plates were incubated at 20°C, until >75% CPE occurred, ∼72 h post infection (hpi). Virus containing media and attached cells was collected and subjected to one freeze-thaw cycle before 30 min 4,000 rpm centrifugal removal of debris at 4°C. Supernatant from each isolate was clarified using 0.22 µm syringe-tip filter and viral particles purified through a 25% (w/v) sucrose pad upon 3 hr 25,000 rpm ultra-centrifugation at 4°C. Virus-containing pellets were re-suspended overnight at 4°C in PBS. Virus stocks were titered by 1:10 serial dilution using confluent EPC cells, divided into 100 µL aliquots, and stored at -80°C. Virus-induced CPE were quantified with a sulforhodamine B (SRB) assay [43].

Infected EPC cells were sampled respectively at 0, 18, 48, 72, or 96 hours post infection (hpi) at a multiplicity of infection (MOI)=1.0 with C03MU or Cell16-a, -b, and -c, and stored at - 80°C. A viral yield assay compared the replication ability of the three test isolates of E16GSa to C03MU virus, by titering media from each time point in 1:10 serial dilutions on BF-2 cells. Plaques were counted 96 hpi and final viral concentration in plaque forming units per mL (pfu/mL) calculated per time point. Antiviral assays followed Ke et al. [43]. UV-irradiated media from each harvested time points was added to EPC cells in 1:3 serial dilutions for 24 h. Cells then were challenged with sucrose-purified C03MU VHSV for 96 h, fixed, and stained with crystal violet (Sigma-Aldrich). CPE plaques were counted and normalized to counts obtained from untreated and uninfected wells. One unit of IFN was defined as the dilution that conferred 50% protection from viral CPE, which value was used to measure the IFN activity (mL) in the testing culture media per time point.

Total RNA was isolated from infected EPC cells following the TRIzol protocol of Gorgoglione [57]. One µg of each RNA sample was reverse transcribed into cDNA by 10 min incubation at 70°C with 100 ng of random hexamer primer (Thermo-Fisher Scientific) and water, for 7 µL total volume. Reactions were cooled to 4°C before adding 13 µL of Moloney Murine Leukemia Virus Reverse Transcriptase (M-MLV-RT) mixture [10X First Stand buffer, 10 mM dNTPs, 0.05 mM random hexamers, 25 U/µL RNasin Plus (Promega), and 200 U/µL M-MLV (Promega)], incubated at 42°C for 1 hr. cDNA was diluted 10 fold in water and archived at - 80°C. 1 µL of each cDNA was tested using RT-qPCR, added to 5 µL Radiant Green Lo-ROX 2× qPCR kit (Alkali Scientific), 50 ng of each oligonucleotide, and water to total 10 µL. Primer were: VHSV-Nse/as, EPC IFN se/as, and β-actin se/as [43]. Reactions and data collection were performed on a C1000 Real Time Thermocycler (Bio-Rad), with initial 3 min 95°C denaturation, 40 cycles of 15 s at 95°C, and 30 s elongation at 60°C. Readings were recorded at the end of each elongation cycle, and threshold values obtained from an automated single point threshold within the log-linear range. Detection of VHSV-N and EPC IFN was normalized to EPC β-actin detection, and normalized to the gene expression of uninfected cell samples. Relative gene expression used the 2^-ΔΔCT^ method [76].

## Results

### Genomic and genic changes

Our analyses of 43 VHSV-IVb whole genome sequences (11,083 NT), plus an additional five from GenBank (GQ385941, KY359355–57), showed no insertions or deletions (Table 1). Of these, 39 sequences were unique (0.81), differing ≥1 NT from C03MU (GenBank: MK782981– MK783014). Isolates C06NP, C06RB, C06SR, C06YP, C06FD, M08AM, C08LEa, C08LEb, and C09MU were identical, designated here as the “C06NP group” (MK782990). No new open reading frames (ORFs) were detected. *G*-gene structures all were identified as glycoproteins.

The 39 IVb gene sequences contained 253 single nucleotide polymorphisms (SNPs) (Table 3), with 85 (0.36) being nonsynonymous, encoding different amino acids (AA). Most SNPs and AA changes occurred in *L* (112 SNPs, 38 AA) and the least *Nv* (14 SNPs, 7 AA), as anticipated from their respective lengths (*L*: 5955 NTs, *Nv*: 369). The non-coding region (NCDS), which does not encode amino acids, contained the highest proportion of SNPs (32 SNPs, 0.043 of the region), whereas *P* and *L* had the lowest (0.019). *Nv* encoded the highest proportion of AA changes (0.057) and *P* the least (0.014). The highest proportion of nonsynonymous: synonymous changes (dN/dS) occurred in *M* (0.53) and the lowest in *P* (0.23). The overall dN/dS for the complete sequence was 0.166.

**Table 3.**
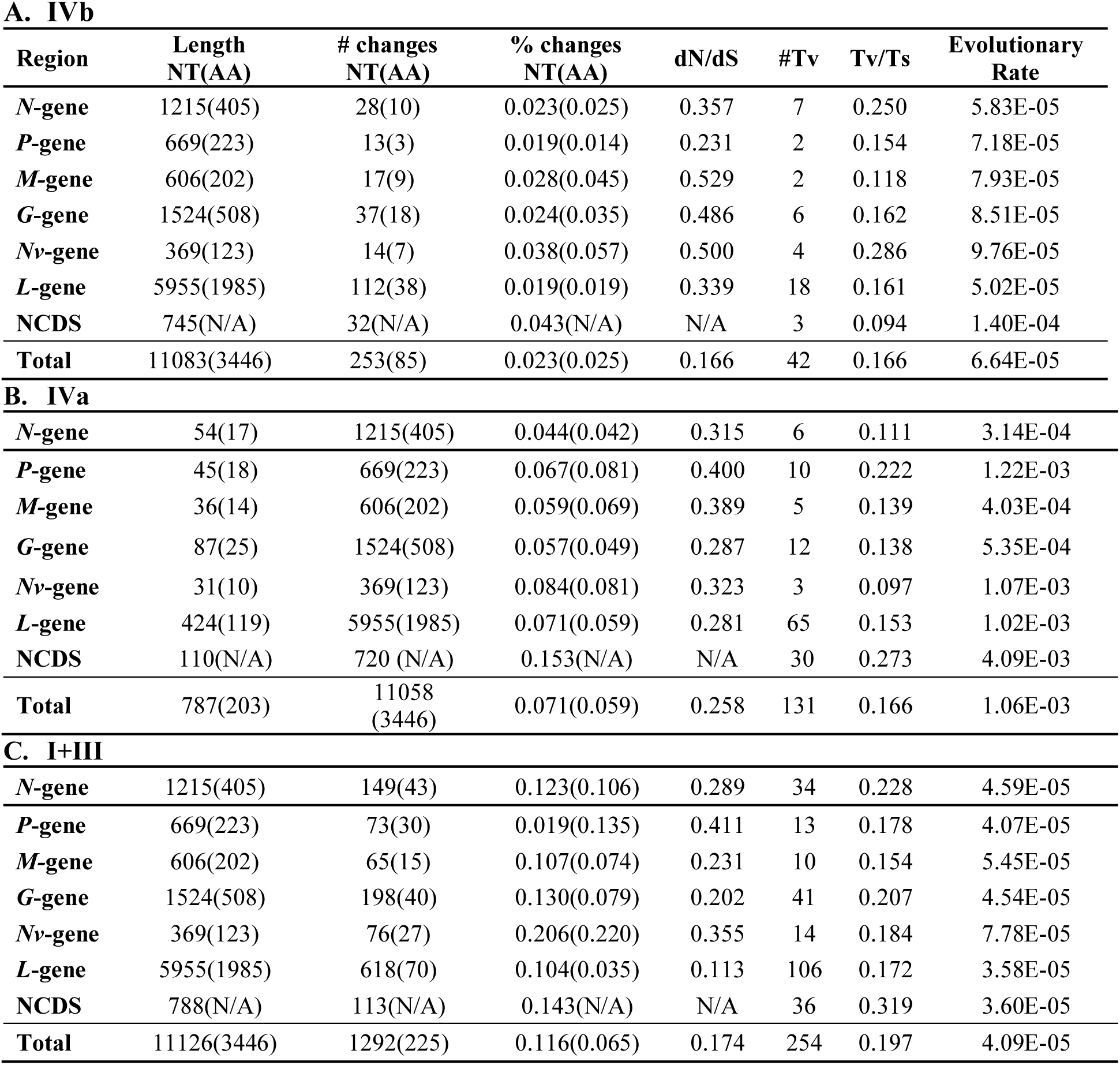
Single nucleotide polymorphisms (SNPs) from each gene’s coding region and the combined non-coding regions (NCDS) in IVb sequence variants of (A) VHSV-IVb, (B) IVa, and (C) Ia, Ib, and III combined. The number of nucleotides (NT) is reported in front of the number of amino acid (AA) changes (the latter are in parentheses). The proportion of nonsynonymous (dN) to synonymous (dS) changes, number of transversions (Tv), and the proportion of transversions to transitions (Ts) are given, along with the evolutionary rate. Totals are in the final row.

**A. IVb**
| Region | Length<br>NT(AA) | # changes<br>NT(AA) | % changes<br>NT(AA) | dN/dS | #Tv | Tv/Ts | Evolutionary<br>Rate |
| --- | --- | --- | --- | --- | --- | --- | --- |
| *N-gene* | 1215(405) | 28(10) | 0.023(0.025) | 0.357 | 7 | 0.250 | 5.83E-05 |
| *P-gene* | 669(223) | 13(3) | 0.019(0.014) | 0.231 | 2 | 0.154 | 7.18E-05 |
| *M-gene* | 606(202) | 17(9) | 0.028(0.045) | 0.529 | 2 | 0.118 | 7.93E-05 |
| *G-gene* | 1524(508) | 37(18) | 0.024(0.035) | 0.486 | 6 | 0.162 | 8.51E-05 |
| *Nv-gene* | 369(123) | 14(7) | 0.038(0.057) | 0.500 | 4 | 0.286 | 9.76E-05 |
| *L-gene* | 5955(1985) | 112(38) | 0.019(0.019) | 0.339 | 18 | 0.161 | 5.02E-05 |
| NCDS | 745(N/A) | 32(N/A) | 0.043(N/A) | N/A | 3 | 0.094 | 1.40E-04 |
| **Total** | 11083(3446) | 253(85) | 0.023(0.025) | 0.166 | 42 | 0.166 | 6.64E-05 |
| **B. IVa** |  |  |  |  |  |  |  |
| *N-gene* | 54(17) | 1215(405) | 0.044(0.042) | 0.315 | 6 | 0.111 | 3.14E-04 |
| *P-gene* | 45(18) | 669(223) | 0.067(0.081) | 0.400 | 10 | 0.222 | 1.22E-03 |
| *M-gene* | 36(14) | 606(202) | 0.059(0.069) | 0.389 | 5 | 0.139 | 4.03E-04 |
| *G-gene* | 87(25) | 1524(508) | 0.057(0.049) | 0.287 | 12 | 0.138 | 5.35E-04 |
| *Nv-gene* | 31(10) | 369(123) | 0.084(0.081) | 0.323 | 3 | 0.097 | 1.07E-03 |
| *L-gene* | 424(119) | 5955(1985) | 0.071(0.059) | 0.281 | 65 | 0.153 | 1.02E-03 |
| NCDS | 110(N/A) | 720 (N/A) | 0.153(N/A) | N/A | 30 | 0.273 | 4.09E-03 |
| **Total** | 787(203) | 11058<br>(3446) | 0.071(0.059) | 0.258 | 131 | 0.166 | 1.06E-03 |
| **C. I+III** |  |  |  |  |  |  |  |
| *N-gene* | 1215(405) | 149(43) | 0.123(0.106) | 0.289 | 34 | 0.228 | 4.59E-05 |
| *P-gene* | 669(223) | 73(30) | 0.019(0.135) | 0.411 | 13 | 0.178 | 4.07E-05 |
| *M-gene* | 606(202) | 65(15) | 0.107(0.074) | 0.231 | 10 | 0.154 | 5.45E-05 |
| *G-gene* | 1524(508) | 198(40) | 0.130(0.079) | 0.202 | 41 | 0.207 | 4.54E-05 |
| *Nv-gene* | 369(123) | 76(27) | 0.206(0.220) | 0.355 | 14 | 0.184 | 7.78E-05 |
| *L-gene* | 5955(1985) | 618(70) | 0.104(0.035) | 0.113 | 106 | 0.172 | 3.58E-05 |
| NCDS | 788(N/A) | 113(N/A) | 0.143(N/A) | N/A | 36 | 0.319 | 3.60E-05 |
| **Total** | 11126(3446) | 1292(225) | 0.116(0.065) | 0.174 | 254 | 0.197 | 4.09E-05 |

In comparison, subgenogroup IVa sequences (Table 3B) possessed 787 SNPs; 203 were nonsynonymous (0.26). Genogroup I/III (Table 3C) contained 1292 SNPs, with 225 AA changes (0.17). Most substitutions for both were in *L* (IVa: 424 SNPs, 119 AAs; I/III: 618 SNPs, 70 AAs). For IVa, *Nv* had the fewest SNPs (31 SNPs, 10AAs), and *M* the fewest for I/III (65 SNPs, 15 AAs). In IVa, the largest proportion of SNPs occurred in the NCDS, and *Nv* and *P* had the most AA changes (0.081). For I/III, *Nv* contained the largest proportions of SNPs (0.206) and AA changes (0.220). The lowest SNP proportions occurred in *N* (0.044) for IVa, and *P* (0.019) for I/III. The least AA changes were in IVa-*N* (0.042) and I/III-*L* (0.035). For both genogroups, the *P* had the highest dN/dS ratio (IVa: 0.400, I/III: 0.411) and *L* the lowest (IVa: 0.281, I/III: 0.113).

IVb possessed the fewest transversions (Tv: IVb=42, IVa=131, I/III=254), tying with IVa in overall ratio of transversions: transitions (Tv/Ts=0.166). I/III had a larger Tv/Ts (transitions) ratio (0.197). Most Tv occurred in *L* for IVb (N=18, Tv/Ts=0.161), IVa (*N*=65, Tv/Ts=0.153), and I/III (*N*=106, Tv/Ts=0.172). All three groups differed in which genomic regions had the fewest Tv (IVb: *P* and *M, N*=2, Tv/Ts=0.118; IVa: *Nv, N*=3, Tv/Ts=0.097; I/III: *M, N*=10, Tv/Ts=0.154). The largest proportion of Tv/Ts occurred in IVb-*Nv* (Tv=4, Tv/Ts=0.286), and NCDS for both IVa (*N*=0.273, Tv/Ts=0.273) and I/III (*N*=36, Tv/Ts=0.310). The lowest Tv/Ts ratios were: IVb-NCDS (Tv=3, Tv/Ts=0.094), IVa-*Nv* (Tv=3, Tv/Ts=0.097), and I/III-*M* (*N*=10, Tv/Ts=0.154).

Per isolate changes were examined for recent IVb isolates. The average number of SNPs/isolate was 16.1, with 6.1 (0.38) AA changes (Sup Table A). The C06NP group possessed the fewest SNPs, with a single non-synonymous change in *L*. Isolates from 2012–2016 contained the highest SNP proportions (averaging 27.9 SNPs, 9.5 AAs), ranging from 14 (M16RGa, 4 AAs) to 38 (E16GSa, 13 AAs). Isolates recovered from cell culture differed by a few SNPs from their original sequences; CellC03 diverged by four SNPs from C03MU and Cell16a–c were two, six, and eight differences from E16GSa, respectively.

SNP changes are shown in Figure 3. C03MU is centrally located nearby a large cluster of isolates, including the C06NP group. Sequences recovered from 2006–08 Lake Erie outbreaks cluster together, differing by 1–4 SNPs. E12FD radiates from this cluster by 31 additional SNPs. Both Budd Lake isolates (B07BS, PS) form an individual branch, sharing one synonymous SNP in *G*. The Lake Ontario isolates (O06SB, O13GS0) [45] group together, sharing 10 SNPs before diverging by 31 SNPs. Five shared changes were nonsynonymous: one in both *M* and *L* and three in *G*. Lake Michigan isolates appear scattered throughout the network, but remain closer to C03MU, with the 2016 isolates differing by 15–16 NT (eight shared). E16GSa–e and derived cell culture isolates form a distant, smaller cluster, sharing 14 NT changes before E16GSc diverges. The remaining seven isolates shared 19 NT changes before differentiating, with E16GSb as central.

**Figure 3.**
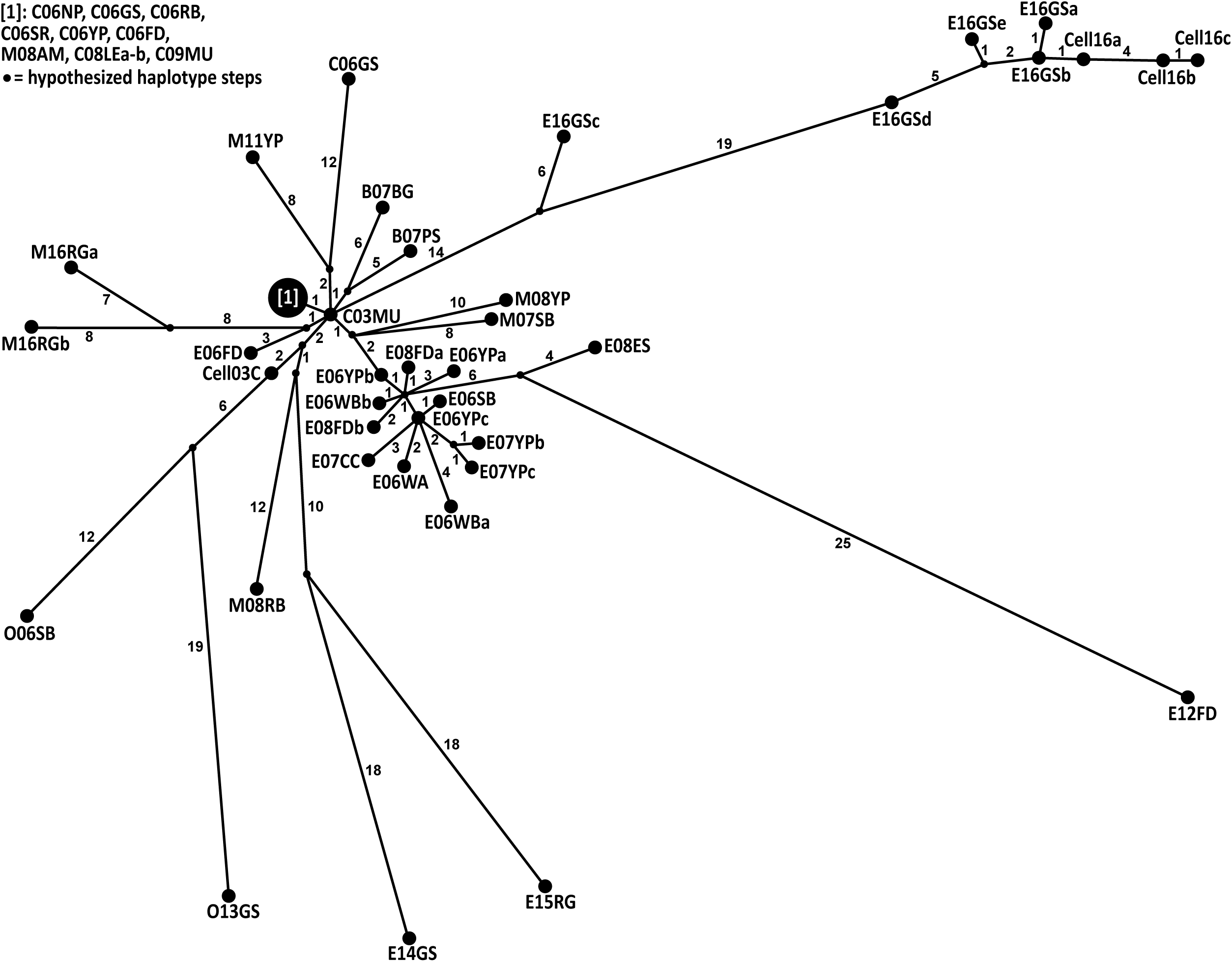
Haplotype network showing genetic relationships among 47 VHSV-IVb full genomes from POPART. Circles sized according to haplotype frequency among isolates. Numbers inside parentheses designate NT differences between each haplotype and the original haplotype, C03MU*. Small, unlabeled black circles = hypothesized haplotype steps.

SNPs distinguished isolates of IVa (Sup. Figure A) and I/III (Sup. Figure B). For the IVa network, the two Japanese isolates were very distant (KRRV9822: 84nt, JFEh001: 253nt) from the nearest Korean isolate (KJ2008). Two identical isolates occurred in I/III, recovered one year apart in Norway (BV06048–52, FA281107). Seven Ib Swedish isolates from 1998–2000 and a 1996 Japanese fish farm isolate form a cluster, differing by 2–22 NT.

The fastest overall evolutionary rate characterized IVa (2.01×10^−3^ substitutions/site/year), followed by IVb (6.64×10^−5^) and I/III (4.09×10^−5^) (Table 3). For IVa, NCDS evolved with the highest rate (7.79×10^−3^), followed by *P* (2.32×10^−3^) and *Nv* (2.03×10^3^). For IVb, NCDS appeared faster (1.40×10^−4^), followed by *Nv* (9.76×10^−5^) and *G* (8.51×10^−5^). The order for I/III was: *Nv* (7.78×10^−5^), *M* (5.45×10^−5^), and *N* (4.59×10^−5^). The slowest regions were IVa: *G* (1.02×10^−3^), *M* (7.68×10^−4^) and *N* (5.97×10^−4^), IVb: *P* (7.18×10^−5^), *N* (5.83×10^−5^), and *L* (5.02×10^−5^), and I/III: *P* (4.07×10^−5^), NCDS (3.60×10^−5^), and *L* (3.58×10^−5^).

Selection pressures for each CDS were examined (Table 4) for IVb, IVa, and I/III. Purifying selection characterized IVb’s *N* (codon 313), *G* (codon 342), and *L* (six codons: 8, 119, 333, 460, 1284, and 1758) genes. For IVa and I/III, FUBAR implicated purifying selection for all genes (Table 4B, C). In IVb, one codon (*L*, 1758) indicated purifying selection and three codons implied diversifying selection – *G* (103, 431) and *Nv* (25). A IVa codon reflected diversifying selection (*G*, 12), along with one in I/III (*N*, 46). MEME implicated diversifying selection for IVb-*G*431 and I/III-*G*477, and three different *L* sites each in IVa (147, 593, 1154) and I/III (112, 474, 1012); none matched.

**Table 4.**
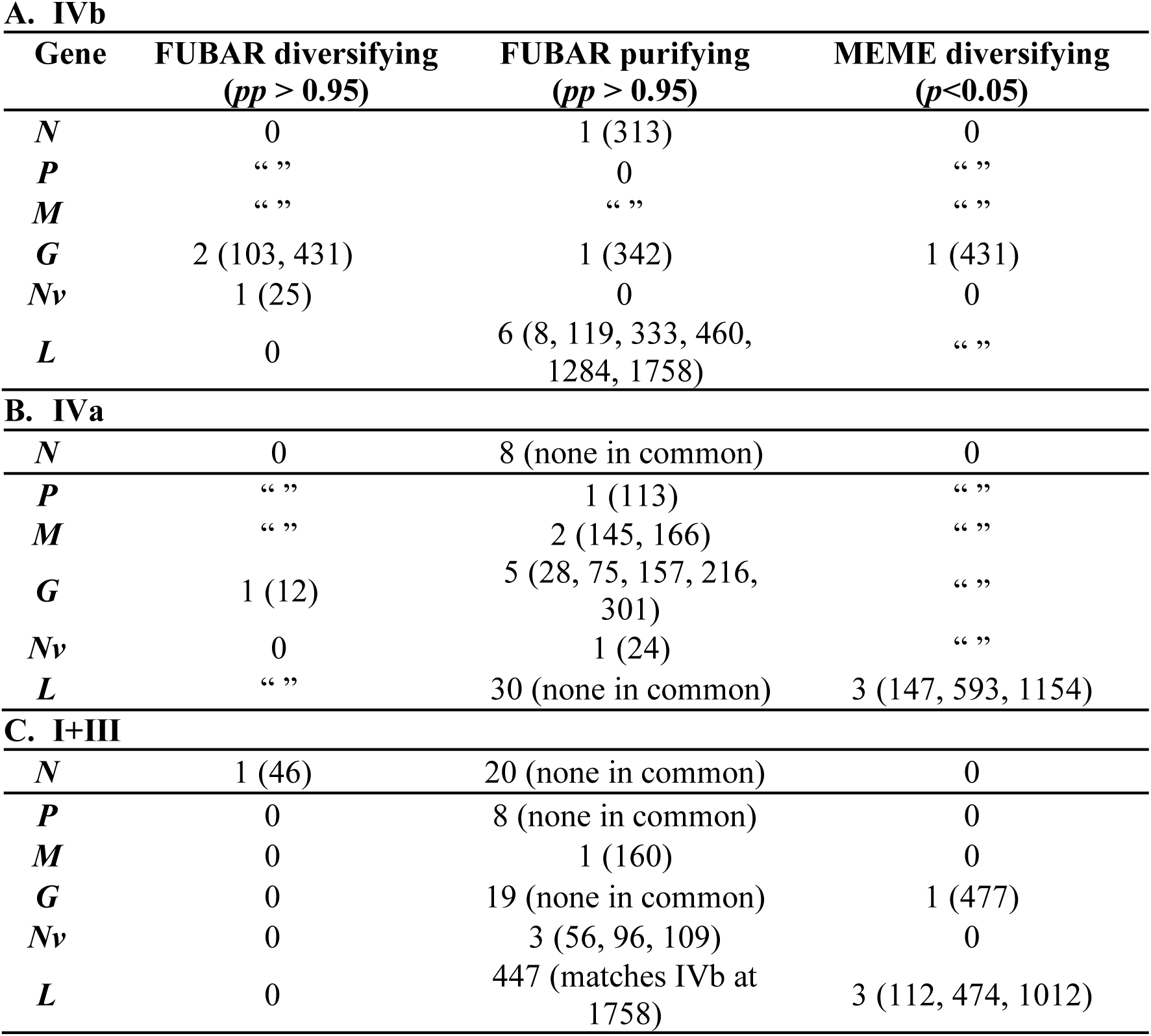
Positive (diversifying) or negative (purifying) selection pressures on individual codons determined by FUBAR (fast, unconstrained Bayesian approximation) and MEME (mixed effects model of evolution) analyses [73, 120] for (A) IVb, (B) IVa, and (C) I/III. None of the codons found in IVa or I/III match the codons under selection for IVb. Results of seven or more codons under selection not displayed. *pp*=posterior probability.

**A. IVb**
| **Gene** | **FUBAR diversifying**<br>**(*pp* > 0.95)** | **FUBAR purifying**<br>**(*pp* > 0.95)** | **MEME diversifying**<br>**(*p*<0.05)** |
| --- | --- | --- | --- |
| *N* | 0 | 1 (313) | 0 |
| *P* | “ ” | 0 | “ ” |
| *M* | “ ” | “ ” | “ ” |
| *G* | 2 (103, 431) | 1 (342) | 1 (431) |
| *Nv* | 1 (25) | 0 | 0 |
| *L* | 0 | 6 (8, 119, 333, 460,<br>1284, 1758) | “ ” |
| **B. IVa** |  |  |  |
| *N* | 0 | 8 (none in common) | 0 |
| *P* | “ ” | 1 (113) | “ ” |
| *M* | “ ” | 2 (145, 166) | “ ” |
| *G* | 1 (12) | 5 (28, 75, 157, 216,<br>301) | “ ” |
| *Nv* | 0 | 1 (24) | “ ” |
| *L* | “ ” | 30 (none in common) | 3 (147, 593, 1154) |
| **C. I+III** |  |  |  |
| *N* | 1 (46) | 20 (none in common) | 0 |
| *P* | 0 | 8 (none in common) | 0 |
| *M* | 0 | 1 (160) | 0 |
| *G* | 0 | 19 (none in common) | 1 (477) |
| *Nv* | 0 | 3 (56, 96, 109) | 0 |
| *L* | 0 | 447 (matches IVb at<br>1758) | 3 (112, 474, 1012) |

### Evolutionary relationships

Significant genetic differentiation characterized IVb over time, indicted by pairwise genetic divergences (Table 5A). The later time group (2012–16) significantly diverged from early (2003–06) and middle (2007–11) groups. Greater divergence was found between the early and later time groups, increasing over time. Population region groupings (Lakes Michigan/Budd, St. Clair, and Erie/Ontario) significantly diverged from each other (Table 5B), with greatest difference between Lakes Michigan/Budd and Erie/Ontario. Significant regional geographic differences occurred between Lakes Erie/Ontario versus other regions for *M* and NCDS, whereas Lakes Michigan/Budd significantly diverged from others with *L*, and *G* differed for Lake St. Clair. *N* differed between Lakes Michigan/Budd and St. Clair.

**Table 5.** Pairwise genetic divergences among VHSV population groupings: (A) sampling time periods, Early (2003–6), Middle (2007–11), and Later (2012–present) and (B) Great Lakes regions (Lake Michigan and Budd Lake, Lake St. Clair, Lake Erie and Lake Ontario) using exact tests (GENEPOP; above diagonal) and *θ*_ST_ divergences (ARLEQUIN; below diagonal). *=*p*≤0.05, **=remained significant (*p*<α) following sequential Bonferroni correction, NS=*p*>0.05.

| **A. Time Groups** |  |  |  | **B. Regional Groups** |  |  |  |
| --- | --- | --- | --- | --- | --- | --- | --- |
|  | **Early**<br>**(N=16)** | **Mid**<br>**(N=16)** | **Late**<br>**(N=11)** |  | **L. St.**<br>**Clair**<br>**(N=10)** | **L. Erie and**<br>**Ontario**<br>**(N=24)** | **L. Michigan**<br>**and Budd**<br>**(N=9)** |
| **Early** | - | NS | ** | **L. St. Clair** | - | ** | ** |
| **Mid** | 0.001 | - | ** | **L. Erie and**<br>**Ontario** | 0.111** | - | ** |
| **Late** | 0.218*<br>* | 0.197*<br>* | - | **L. Michigan**<br>**and Budd** | 0.264** | 0.348** | - |

The *Novirhabdovirus* phylogenetic tree (Figure 4) defines each species with 1.00 posterior probability (*pp*) and 100% bootstrap support (bs). IHNV and HRRV are sister species, comprising the sister group to VHSV+SHRV. The 79 genomic VHSV sequences form two clades: European genogroups I–III and North American/Asian IV. Within I–III, the single VHSV-II isolate is basally located. Two of the five VHSV-III genotypes group with two Ia isolates (Hededam DK35928), thus not monophyletic (i.e., not a clade). In contrast, the seven Ib isolates are a well-defined clade, and the III isolates comprise another. Genogroup IV (Figure 3) possesses 1.00 *pp* and 100% bs support, with IVa (*N*=25) and IVb (*N*=39) on separate branches, having 1.00 *pp* and 100% bs support.

**Figure 4.**
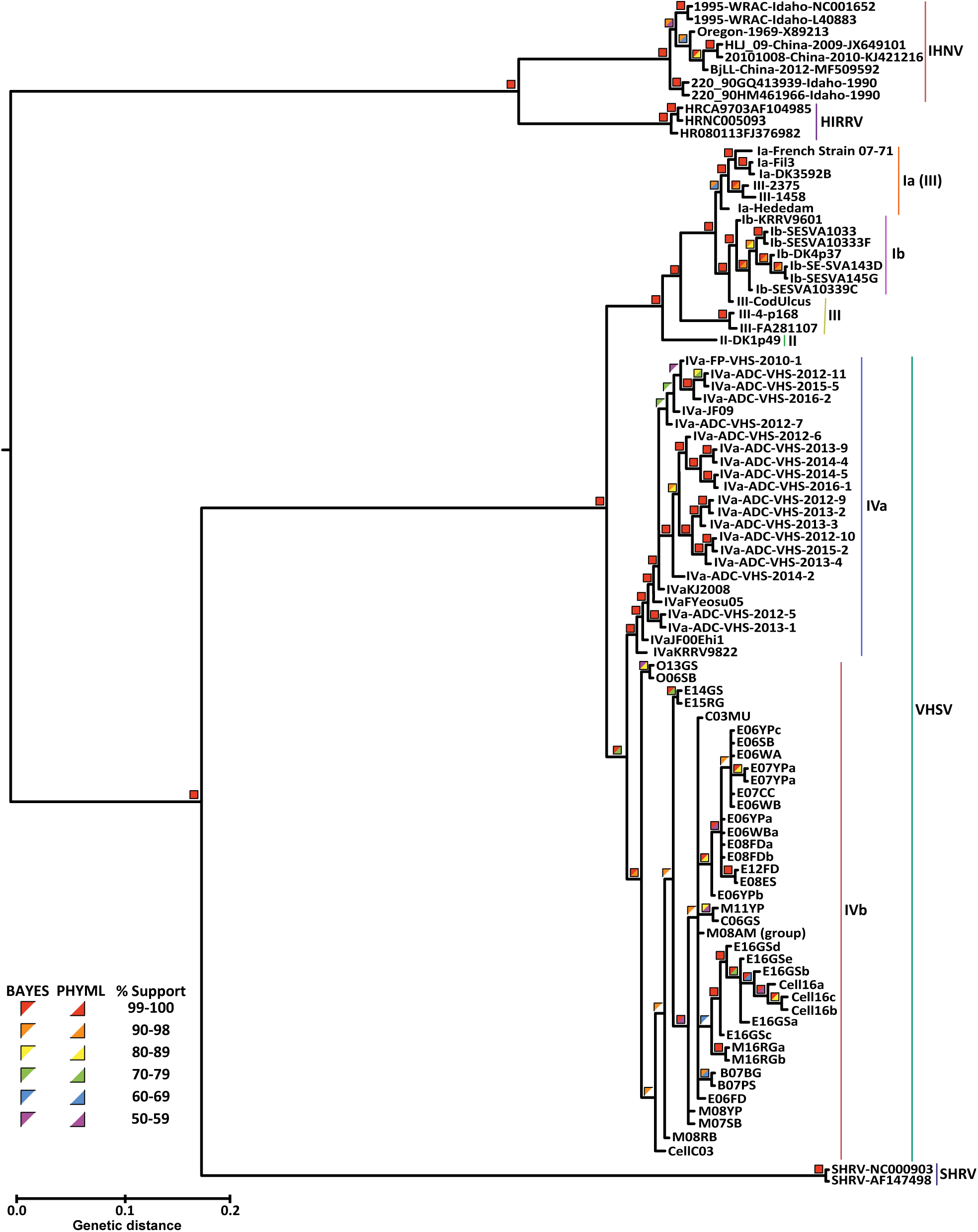
Novirhabdovirus phylogenetic tree, based on full genome sequences (see Tables 3.1 and 3.2), from maximum likelihood and Bayesian analyses. Colored squares designate support values, top left half = Bayesian posterior probabilities, bottom right = 500 bootstrap pseudoreplicates. Tree is rooted to the Snakehead Rhabdovirus (SHRV, GenBank: AF147498).

The IVb phylogenetic tree (Figure 5) contains several inner clades, with M08RB appearing basal (weakly supported), followed by O06SB, CellC03, and O13GS, respectively (0.60–.90 *pp*/<50% bs). Two Lake Erie isolates from 2014–15 (E14GS, E15RG) group together (1.00 *pp*/74% bs), followed by two Lake Michigan 2007–08 isolates, located prior to the main cluster of the remaining 39 isolates, including C03MU. Within the major group are two clades containing two isolates (B07BG/B07PS: 0.80–.89 *pp*/66% bs; M11YP/C06GS: 0.80–.89 *pp*/<50% bs) and two larger clades. The first larger clade incorporates the 2016 isolates (0.60– .69 *pp*/<50% bs), further subdivided in two, separating Lake Michigan (1.00 *pp*/98% bs) from Lake Erie and cell culture derivatives (1.00 *pp*/100% bs). A second well-defined clade (1.00 *pp*/83% bs) contains 14 isolates, including most early to middle (2006–08) Lake Erie genotypes, along with its very distinct 2012 isolate. The 2012 genotype is the most distant, comprising the sister taxon to E08ES (1.00/94%), which are linked by four shared SNPs and 2 AAs (*M*: one SNP, one AA; *L*: two SNPs, one AA; NCDS: one SNP). E12FD and E08ES share an additional five SNPs: two in *N* (two AA), and two in *L* (one AA).

**Figure 5.**
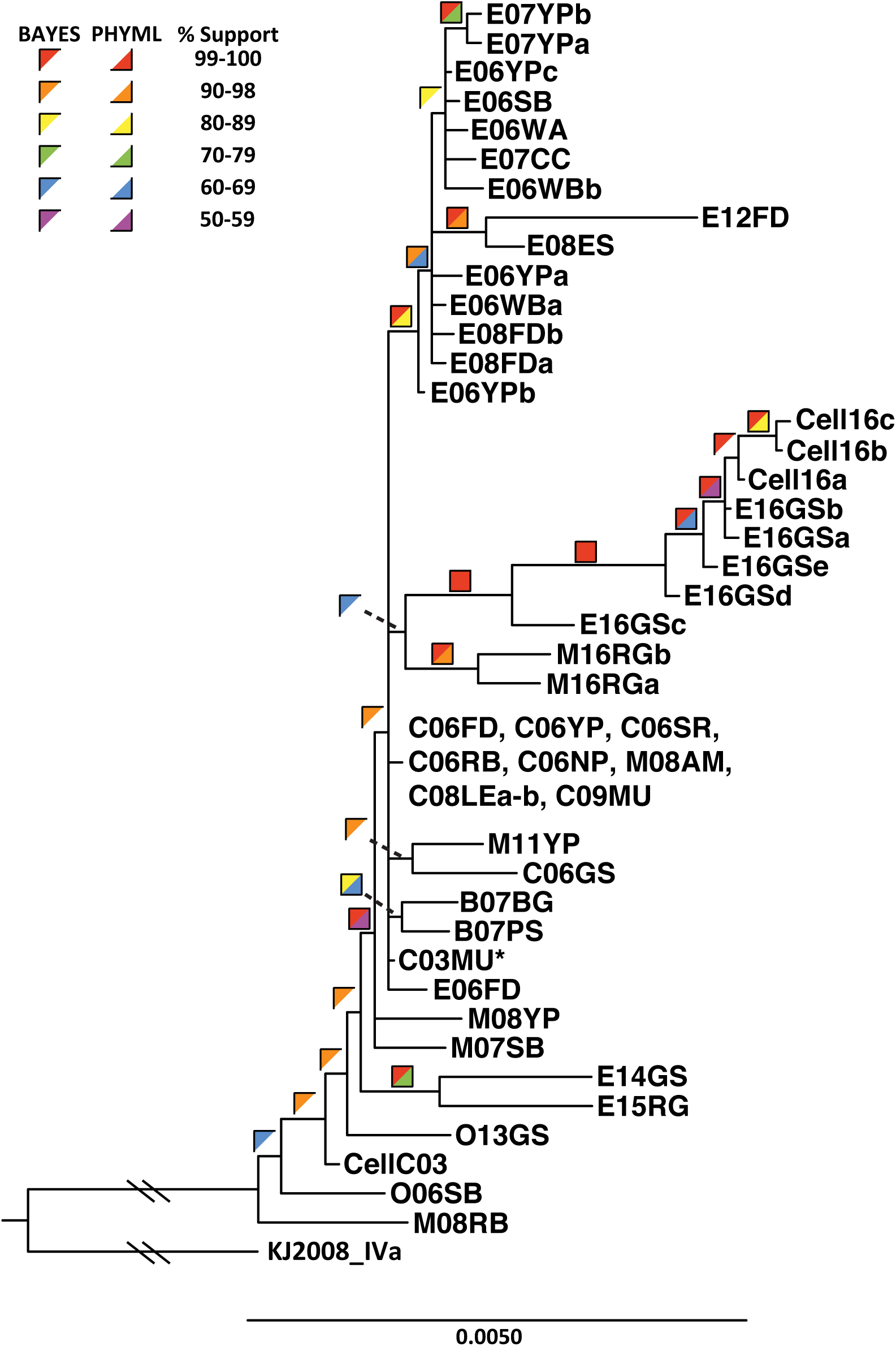
VHSV-IVb phylogenetic tree of IVb whole genome haplotypes, with maximum likelihood and Bayesian analyses. Colored squares = support values, top left half = Bayesian posterior probabilities, bottom right half = 1450 bootstrap pseudoreplicates. Hashes represent cropped region for visualization. *=original IVb isolate. The tree is rooted to substrain IVa (GenBank: JF792424).

BLAST searches returned no similar sequences to *Nv* when novirhabdoviruses were omitted. The NT phylogeny (Figure 6A) resolved all genes as comprising distinct clades, except for *Nv*. The phylogeny contained two groups of clades: *M*+*G* and *N*+*P*+*L*. SHRV-*Nv* grouped with *G*. IHNV-*Nv* and HIRRV-*Nv* are sister groups, sharing a common ancestor with the *N* clade. VHSV’s *Nv* gene appeared a divergent part of the *L* clade. All *Nv* clades were fully supported by Bayesian analysis.

**Figure 6.**
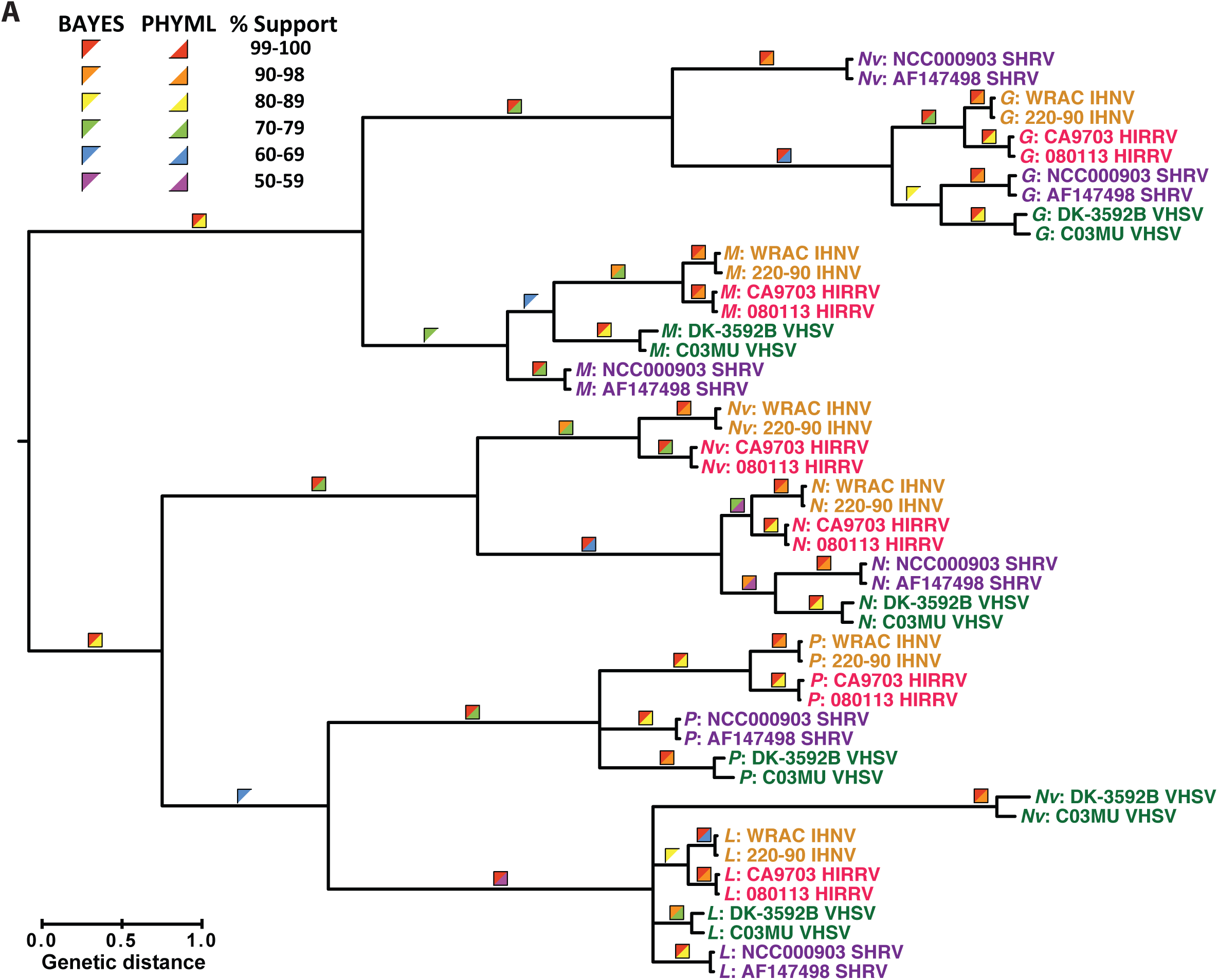
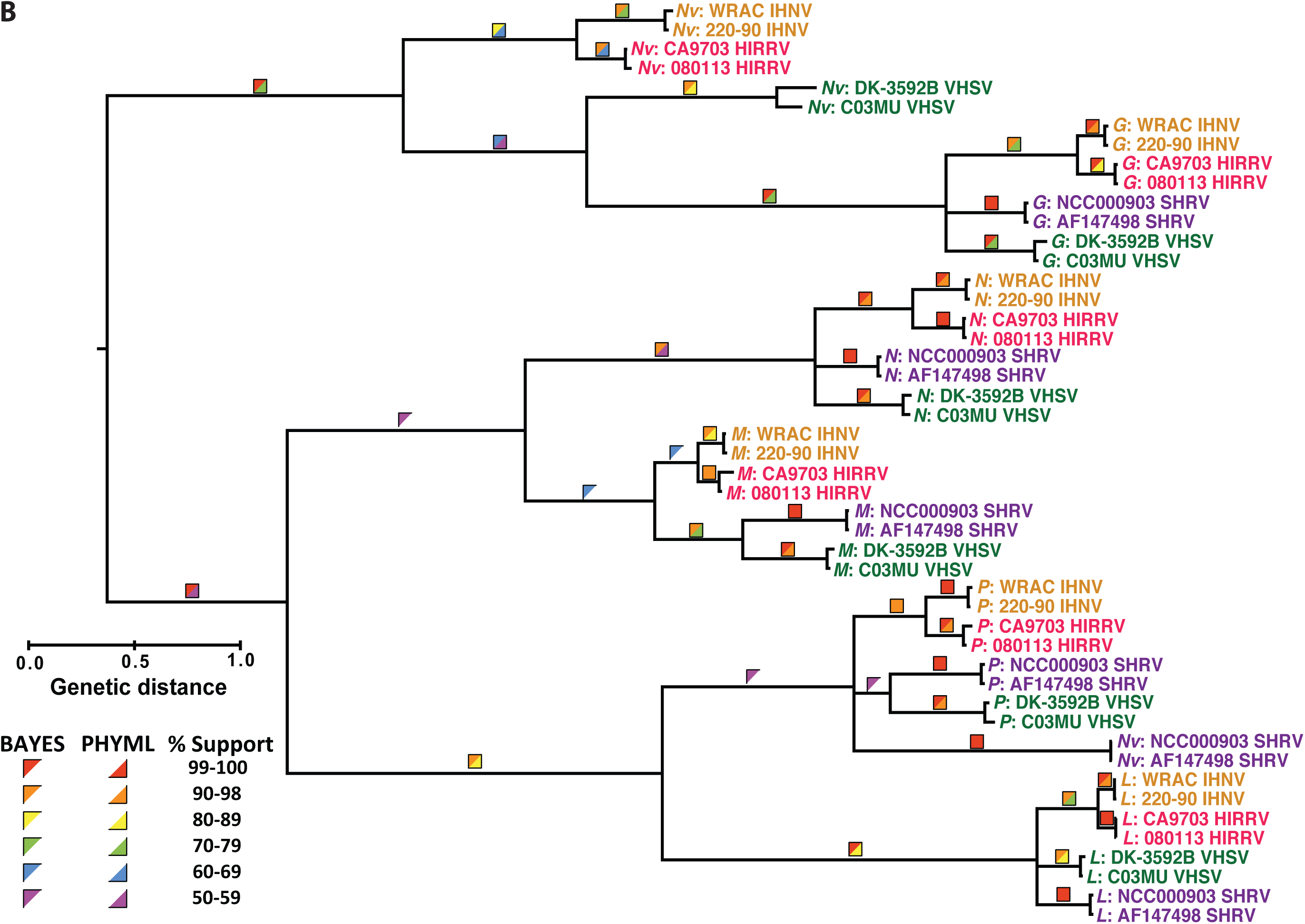
Phylogenies of individual novirhabdovirus gene, based on full coding sequences of two representatives sequences of all species (see Tables 3.1 and 3.2), with maximum likelihood and Bayesian analyses for (A) NT and (B) AA. Colored squares designate support values, top left half = Bayesian posterior probabilities, bottom right=2000 bootstrap pseudoreplicates. Tree is rooted to the Snakehead Rhabdovirus (SHRV *L*-gene, GenBank: AF147498).

When AAs alone were considered (Figure 6B), the IHNV and HIRRV-*Nv* clade was sister to VHSV-*Nv* and the *G* clade. SHRV-*Nv* formed its own clade, more closely related to the *P* clade, but was not well supported.

### Differences in cytopathicity and immune response

A series of cell-based studies were used to determine whether sequence variations impacted viral function: SRB assays to assess cytopathicity, virus yield assays to estimate viral production, antiviral assays to assess IFN production, and qPCR to determine levels of IFN and VHSV-IVb mRNA production. Sequencing confirmed each isolate’s identity and revealed changes acquired during cell culture propagation. The cell culture amplified control, CellC03, differed from C03MU by four NTs and two AAs, *M* (2312, NT:C–T, AA:T–I) and *G* (3397, NT:G–A, AA:K–D; 4007, NT:C–G, AA:G–G; 4394 NT:G–A, AA:V–V). Cell16a–c differed by 34–38 SNPs (13–14aa) from C03MU (Sup Table A). From E16GSa, Cell16a differed by two NTs: *N* (737, NT:C–A, AA:T–T) and *G* (4117, NT:G–A AA:D–N), Cell16b differed by four additional changes (six total) in *L* (6962, NT:C–A, AA:A–E; 7038, NT:G–A, AA:E–E; 7047, NT:T–C, AA:C–C; 7647, NT:T–C, AA:H–H), and Cell16c had one further NT change (one total) in *L* (6456, NT:C–T, AA:L–L).

SRB staining examined CPE elicited at different MOIs across the four isolates. The 2016 isolates exhibited less CPE than CellC03 when tested at MOIs >1×10^−3^ (Figure 7). Although not statistically different across all isolates and at all dilutions, the 2016 viral isolates tested were generally less cytotoxic than CellC03. Consistent with this observation, viral yield assays demonstrated that at 96 hpi, CellC03 produced significantly more virus than the 2016 isolates (Figure 8), yielding nearly 100-fold more infectious particles as compared to Cell16a–c.

**Figure 7.**
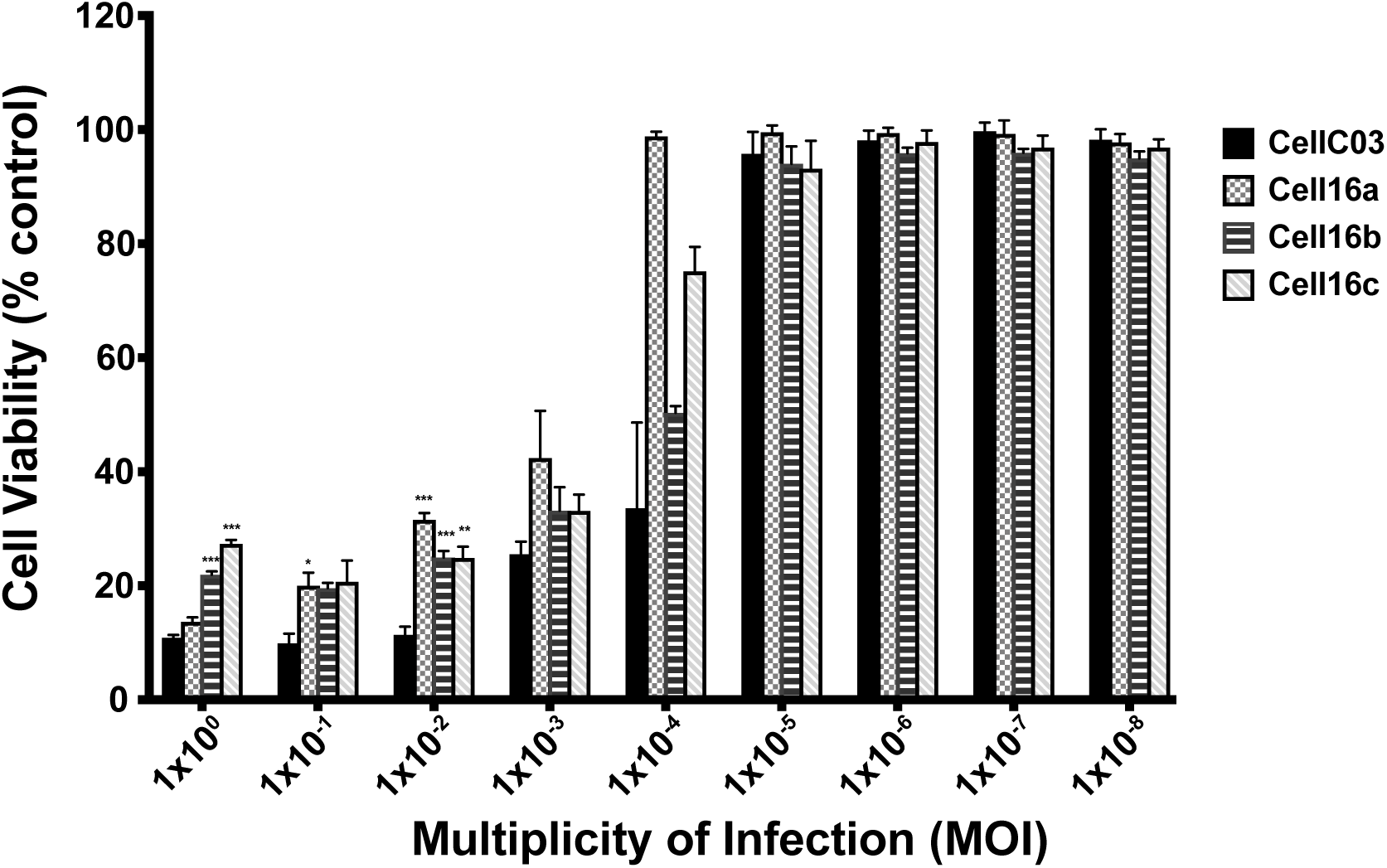
Host cell viability as measured by Sulforhodamine B (SRB) assay in EPC cells infected with MOIs (MOI=1×10^−8^–1.0) of four VHSV-IVb isolates, 96 hpi. Average values from a single experiment (conducted in triplicate) are shown, and are representative of at least three independent experimental replicates. Standard error bars, \**p*<0.05; \*\**p*<0.01; \*\*\**p*<0.001.

**Figure 8.**
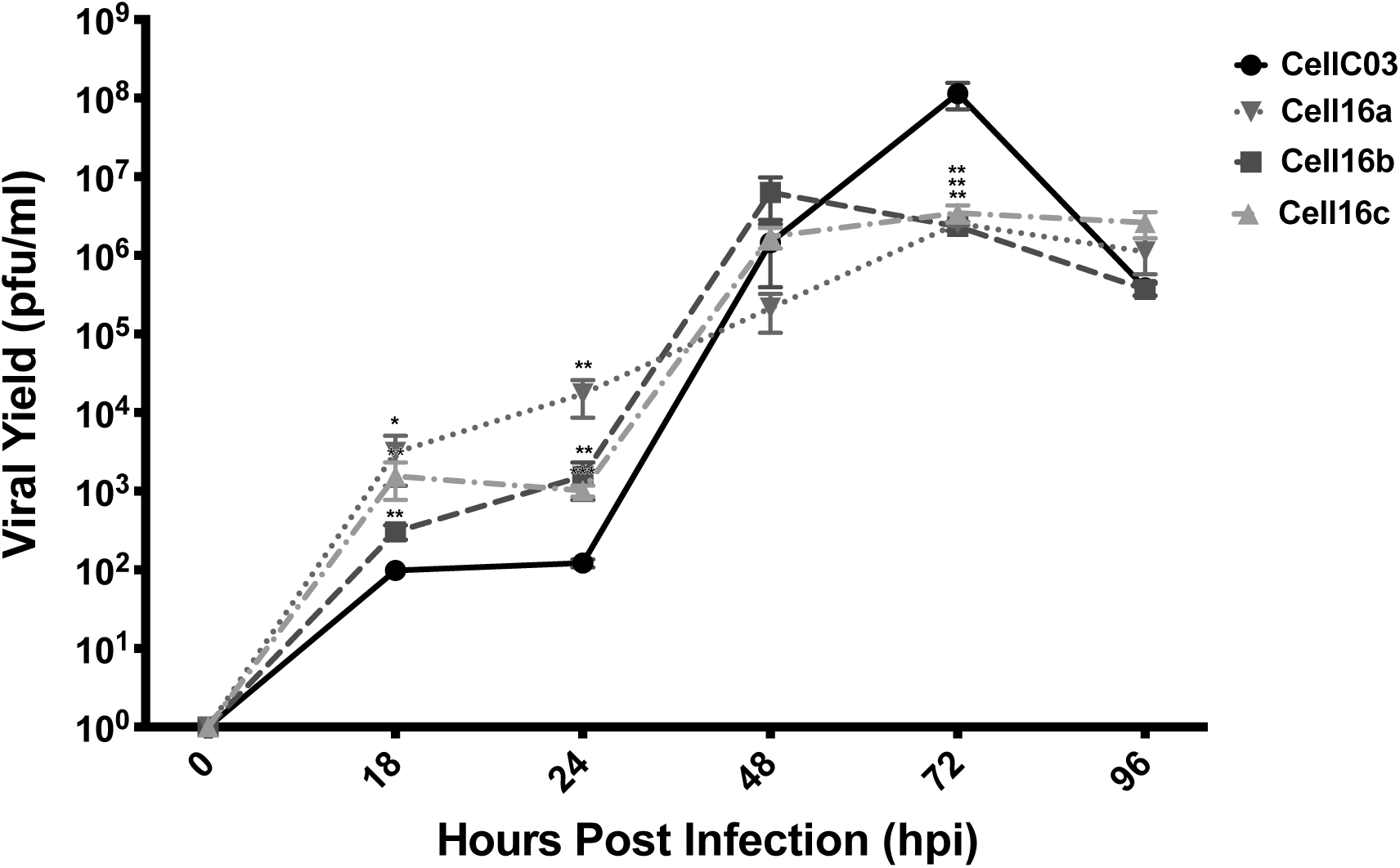
Viral yield assay comparison of infectious viral particles produced (pfu/ml) in wild type (CellC03) and three 2016 VHSV-IVb samples, in BF2 cells 96 hpi, following exposure to media from collection time post infection. Mean values from a single experiment (conducted in triplicate) are shown, and are representative of at least three independent experimental replicates. Standard error bars are shown; \**p*< 0.05; \*\**p*<0.01; \*\*\**p*<0.001.

### IFN production and expression of VHSV and IFN mRNAS

IFN production was measured in an antiviral bioassay, which revealed earlier onset of antiviral immune response in cells infected with Cell16a, yielding significantly more IFN at 24 hpi than CellC03 (Figure 9). All isolates produced comparable IFN activities after 24 hpi (Figure 9). Since IFN antiviral bioassays are relatively insensitive to minimal changes, we extended the previous studies by assessing IFN mRNA synthesis in infected cells with RT-qPCR. IFN mRNA was expressed significantly more in Cell16a–c, compared with the CellC03 control across all time points beyond 24 hpi (Figure 10). These data suggest significant difference in ability of the 2016 isolates to induce cellular IFN expression, or significant reduction in their suppression of IFN expression. Consequently, by 48 hpi, CellC03 produced the highest overall amount of viral RNA at 48 hpi, which continued until 96 hpi, by which point cell death in all cultures led to reduced viral expression. All 2016 isolates produced similar amounts of viral RNA.

**Figure 9.**
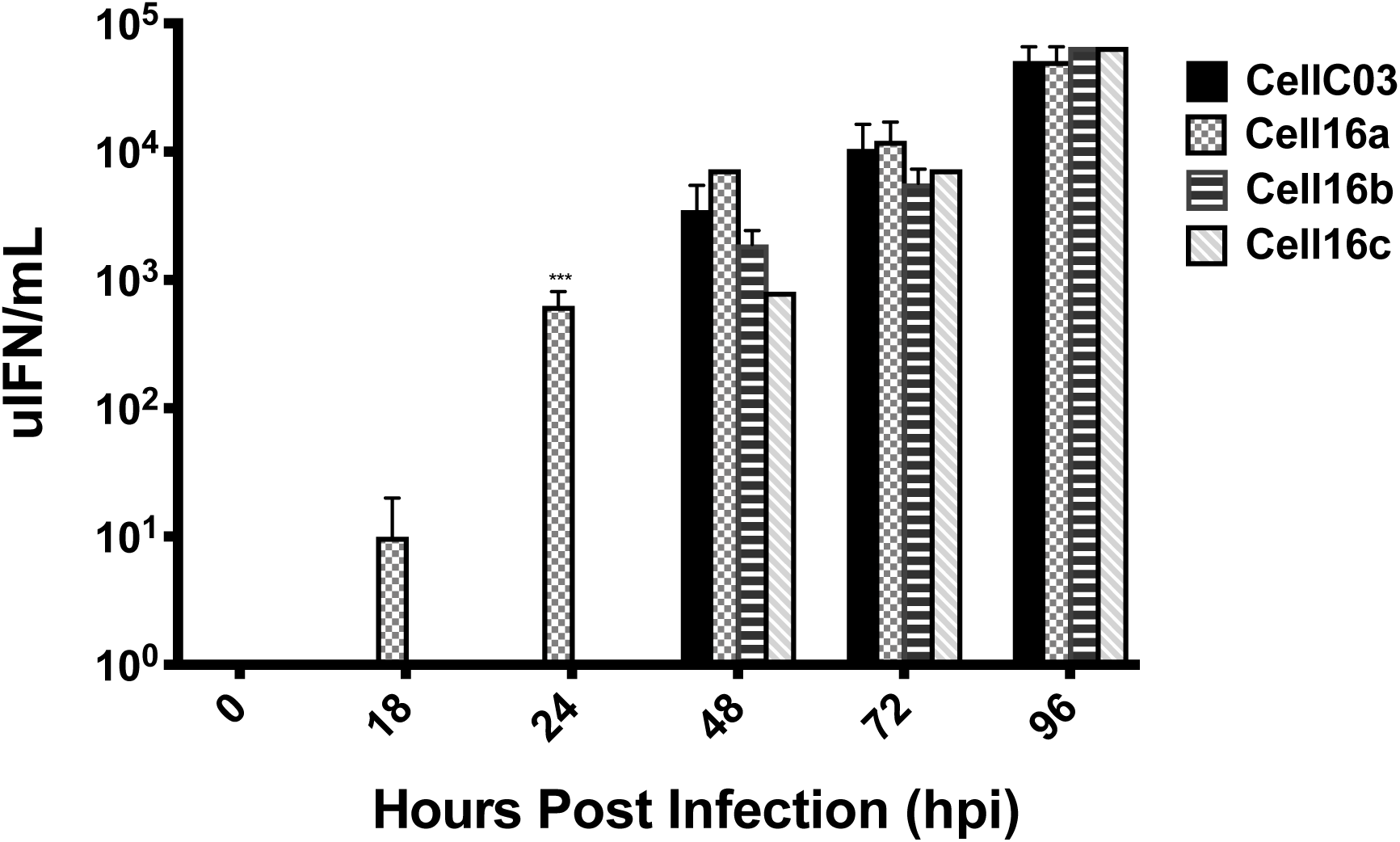
Antiviral assay comparison of host IFN suppression between reference (CellC03) and three more recent VHSV-IVb isolates, in EPC cells 96 hpi following exposure to UV irradiated media collected at the above time points. Values are quantified as the number of antiviral units (uIFN) per mL. Mean values from a single experiment (conducted in triplicate) are shown, and are representative of at least three independent experimental replicates. Standard error bars, \**p*< 0.05; \*\**p*<0.01; \*\*\**p*< 0.001.

**Figure 10.**
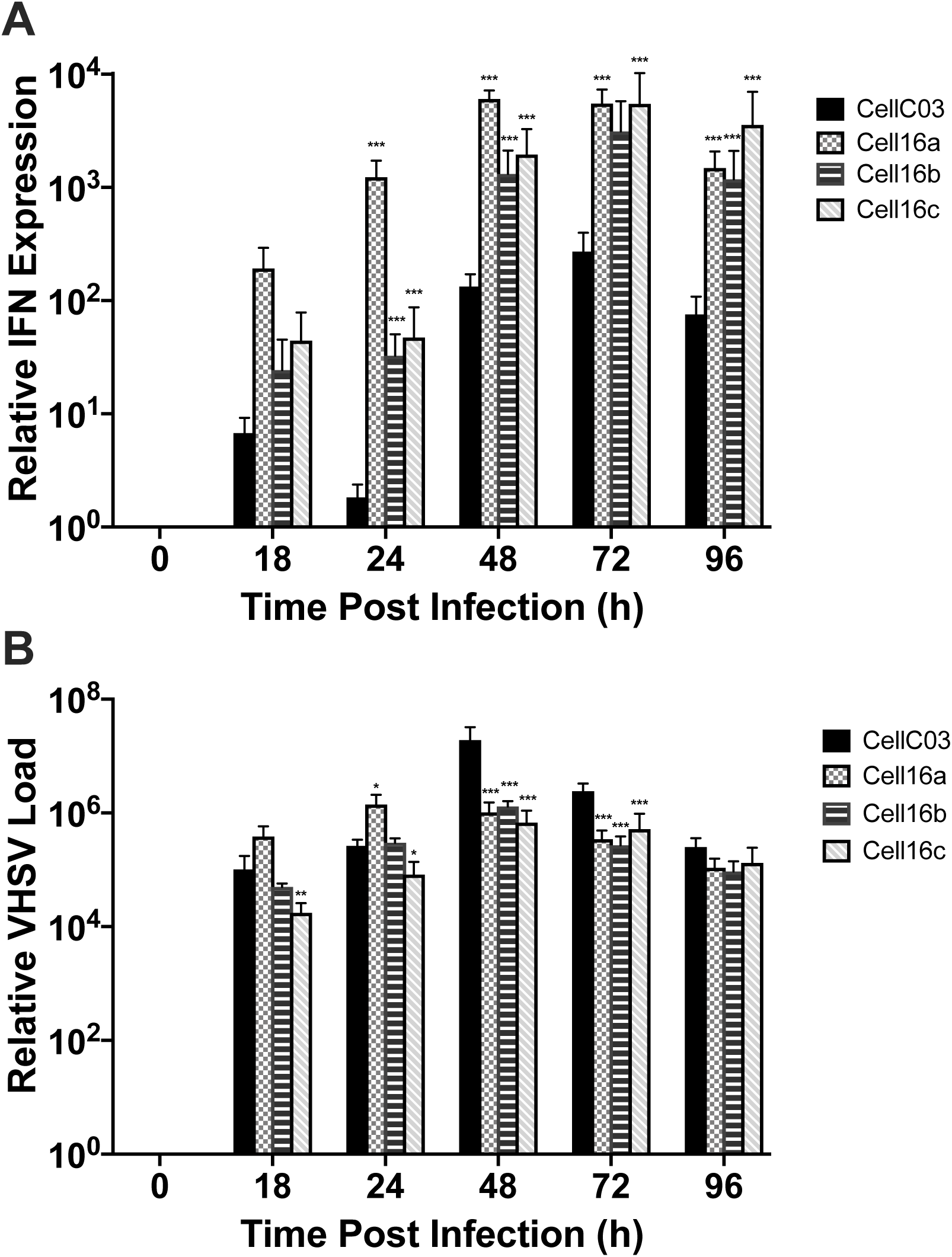
Gene expression of host immune response and viral RNA produced. qPCR comparisons between reference (CellC03) and three 2016 VHSV-IVb isolates for (A) EPC IFN and (B) virus detected for samples at each of the above time points. Data were normalized to β-actin mRNA levels. Mean Average values from a single experiment (conducted in triplicate) are shown, and are representative of at least three independent experimental replicates. Significance was calculated from Ct values and transformed values (2^-ΔΔCT^) are shown. Standard error bars, \**p*< 0.05; \*\**p*<0.01; \*\*\**p*< 0.001.

## Discussion

### Evolutionary patterns

We analyzed 46 sequences of VHSV-IVb for the largest Novirhabdovirus genomic investigation. The greatest proportion of SNPs occurred in the NCDS (4.3%) and the *Nv*-gene (3.8%). In IVb-*Nv*, none of the isolates we sampled had >two SNPs, suggesting that sequence conservation is important. An experimental study on *Nv* function indicated that mutations at codon positions 36, 39, and 41 exhibited greater host immune suppression [26]; our sequences lacked SNPs at those locations. Since *Nv* is not essential for viral replication [19], its SNP variations may be more tolerable for the virus. Large proportions of SNPs likewise characterized NCDS in VHS-IVa and I/III, illustrating sequence flexibility. There is no evidence that NCDS regions are transcribed or play a functional role in any rhabdovirus [2], thus these SNPs likely are selectively neutral.

Diversifying selection was indicated for two IVb genes: *G* and *Nv*. Unlike our findings, a study of just four IVb genome sequences (C03MU, O06SB, O13GS, and E14GS) by Getchell et al. [45] did not uncover diversifying selection for either *G* or *Nv*. Getchell et al. [45] discerned variation in two *N-*gene codons and one codon each for *M* and *L*, which were unsupported in our investigation. We found evidence of purifying selection acting on single *N-* and *G-*gene codons, and on six codons in *L*. In our IV analyses. Thus, fewer codons were under purifying or diversifying selection than in Getchell et al.’s [45] findings. None matched between the two analyses, with ours being the larger, more robust dataset.

Purifying selection was the major evolutionary force discerned across all VHSV genogroups, subgenogroups, and samples. Our study identified ≥one codon per gene under negative selection, for IVa and I/III. No NT sites under selection were in common between IVb and the other genogroups, with one exception: codon 1758 in *L* for IVb and I/III. Our dN/dS ratio further indicated purifying selection, with values being >0.5 across all genes and VHSV genogroups. Other investigations found that purifying selection regulates VHSV evolution [30, 77], as well as in other rhabdoviruses [78] and RNA viruses in general [79]. He et al. [77] compared dN/dS ratios for individual genes among VHSV genogroups, with all six genes having lower dN/dS than found with our data. However, most of the IVb (N=45) and IVa (*N*=18, from [80]) sequences used in our analyses were unavailable to He et al. [77].

Positive selection was indicated for changes in three *G*-gene codons: two in IVb and one in IVa. Positive selection on viral genes constitutes evidence that supports the host-pathogen “arms race”, which typically suppresses host immune responses [51]. Abbadi et al. [30] examined 108 VHSV-Ia full-genome *G* isolates, discerning positive selection on two codons (258, 259). He et al. [77] also elucidated positive selection for I at codons 258 and 476, but not for 259. Our codon results did not match either of those studies.

The host immune system exerts strong selective pressure on viral mutation rates. Host suppression by multifunctional viral proteins is a critical determinant of success for all RNA viruses [51], including Human Immunodeficiency Virus (HIV) [81]. Viral RNAs can suppress host RNA interference pathways involved in antiviral immunity [82] or block the abilities of host antiviral proteins to engage with viral dsRNAs and mount an immune response [83].

We found that overall evolutionary rates across the entire genome were slower than were previous estimates based on partial gene sequences [12]. These differences stem from our present use of more samples and complete genome sequences, which included conserved areas that were previously omitted.

Among the three VHSV genogroups, V-IVa exhibited the fastest rate – 2.01×10^−3^. He et al. [77] estimated a rate of 5.60×10^−4^ for 48 IVa *G*-gene sequences, which was slower than our IVb rate for *G* of 1.02×10^−3^. Our inclusion of recently published diverse sequences from Korea between 2012–16 [80] may have increased this, as those isolates differed by >20nt. Just two complete sequences of IVa were from Japan, which were even more different than the Korean isolates. This suggests that the Korean isolates are diverging from the Japanese IVa isolates, but additional IVa genomes from Japan are needed for robust conclusion. IVa first was detected along the North American Pacific Coast [32, 33, 34], but no full-genome sequences then were available. All IVa Korean isolates were obtained from aquaculture; such transportation of infected fishes may introduce VHSV to naïve fishes and different environmental conditions, which can enhance adaptation and dramatically alter the virus, as observed in IHNV [84] and HIRRV [85].

Some Ia, Ib, and III isolates grouped together in our phylogeny, indicating that their previous phylogenetic relationships, based on partial genes [12, 30, 86], had been poorly resolved in the earlier analyses. We calculated that genomes of VHSV genogroups I/III evolved at an overall rate of 4.09×10^−5^, more similar to that of IVb. Previous *G*-gene rate estimates for VHSV-I and -III [77] using fewer complete gene sequences were faster than ours (4.54 × 10^−5^), having been based on 201 I sequences (5.57×10^−4^) and seven III isolates (1.63×10^−3^). Evolution of subgenogroup Ia had been estimated at 1.74×10^−3^ (*N*=34) [23] to 7.3×10^−4^ from 108 *G*-gene Italian isolates [30]. Our combination of the European groups may have yielded our slower rate. Despite these variations, all of the above estimates are within the known range of evolutionary rates for RNA viruses [87](Duffy et al. 2008).

### Phylogenetic patterns: Novirhabdoviruses

The phylogeny from the whole genome (Figure 3) is congruent with prior partial gene studies [12, 14]. Two main sister groups characterize the novirhabdoviruses: IHNV+HIRRV and VHSV+SHRV, with VHSV+SHRV diverging first from the common ancestor. This relationship is congruent with Kurath’s (2012 [18]) analysis of complete *N-*gene sequences. Two studies examined rhabdoviruses using partial *L-*gene sequences [88, 89], which likewise supported our sister group pairing of IHRV+HIRRV. Our phylogenetic consensus tree yielded 100% ML and Bayesian support for these relationships, whereas Kurath [18] found just 76% support for the SHRV+VHSV clade using *N*. Thus, our analysis of the entire genome using a larger number of isolates significantly increased confidence in resolving novirhabdovirus evolutionary relationships.

Phylogenetic results for VHSV relationships from whole genome sequences support most known genogroups and subgenogroups, except that III and Ia are not monophyletic. Prior analyses of *G*-gene sequences by Dale et al. [90] and Ghorani et al. [31] thus were incorrect in separating III and Ia. Genogroup Ia occurs in freshwater hosts [23], whereas both III and Ib are marine [90]. All three are capable of infecting rainbow trout (*Oncorhynchus mykiss*), which spends portions of its lifecycle in both environments. Our results indicate that Ib is monophyletic, whereas III and Ia comprise a single taxon, indicating that they need to be re-defined as a single taxon. Additional III and Ia isolates should be completely sequenced.

### Phylogenetic patterns: VHSV-IVb

Our phylogenetic tree contains several clades of closely related IVb isolates, which correspond to sampling location and year. We resolve an in-depth view of the virus in Lake Erie, where a large portion of our samples were collected. The inner-most branches of the phylogeny suggest that at least two separate infections led to the major outbreaks, since one of the 2006 isolates (E06FD) appeared more closely related to C03MU and the Lake St. Clair outbreak samples.

The differentiated Lake Erie clade from 2006–08 and 2012 suggests its separate evolutionary history from the 2014–15 and 2016 clades. Broad diversification characterized the 2006 outbreak, which continued along that trajectory in 2007–08, with minor variants. The lone 2012 sample, although evolutionarily distant, appears more closely related to the E06–08 clade, than to Lake Erie samples from subsequent years. E12FD forms a sister clade with E08ES, sharing an additional four AA changes in the coding regions of *N, P*, and *L*. These AA changes possibly provided some advantage or lacked deleterious effects, since the substitutions remained conserved for four years. Although most of that clade was recovered from Percidae hosts, several of its isolates were in freshwater drum. In addition to the 2006 and 2008 Lake Erie cases [11], freshwater drum die-offs were reported in 2005 from Lake Ontario [9], which appeared more closely related to E06FD.

Both of our 2007 samples from the inland Budd Lake form a clade, located near C03MU. Wayne County, Michigan, which is adjacent to Lake St. Clair that is a highly urbanized area with the largest number of registered anglers in the state [91]. Anglers traveling from that area to Budd Lake may have unknowingly transported VHSV, perhaps via live bait [12, 14, 92].

The 2014–15 Lake Erie VHSV-IVb genotypes differ from one another, but share a sister relationship branching from the main part of the phylogeny, located closer to C03MU. Genotypes E14GS and E15GS shared the host species of gizzard shad and round goby with the 2016 isolates, but are genetically distant from the latter. The 2016 genotypes form a clade, which is linked by SNPs as shown in the genotype network. Interestingly, these occurred in different host species, with the E16 isolates exclusively recovered from gizzard shad, and both M16 isolates in round goby. Other studies identified potential evolutionary radiation of IVb in the round goby host, particularly in Lake Ontario [40, 75]. There have been relatively few VHS positive round goby samples detected in Lake Michigan, despite its reported deaths during the 2008 outbreaks [16]. Round goby has been implicated in spreading the virus [93], often is used as bait, and is transported among water bodies by anglers [94].

Other VHS-positive isolates were recovered from Lakes Michigan and Erie in 2016, whose viral titers were too low for genome sequencing [75], further indicating that infection levels differ spatially and temporally. One of the first 2006 outbreaks intensely affected the Lake St. Clair gizzard shad population [16], as well as its 2017 outbreak (G. Whelan, personal communication 2017). Gizzard shad populations frequently experience large die-offs from cold weather [95], water temperature fluctuations, and/or spawning activities [96], coinciding with the optimal temperature range of VHSV-IVb (12–18 °C [97]; 10–14 °C [98]. In addition to the 2016 isolates, IVb-positive gizzard shad also occurred in Lakes St. Clair, Erie, and Ontario dating back to 2006; however, only its 2016 isolates were closely related. Possible host specialization should be examined in future investigation.

All but one of our identical sequences (C06FD group) were collected from Lake St. Clair over a five-year span. This consistency suggests that the C06FD genome may have been prevalent and locally adapted to environmental conditions and host species. It differed from C03MU by a single AA in the *L-*gene, and was found in six different fish host species and the sole two known invertebrate host species [99, 100]. Yusuff et al. [101] discerned no effect on host specificity when swapping *L*-genes between C03MU and Ia’s DK-3592B. *L* variants appear to differ in optimal temperature range, since swapping *L* from IVa into IVb led to increased virus at higher temperatures, than occurred for IVa [102]. The single AA change between the C06FD group and C03ML was not shared with IVa; instead C03ML and IVa were identical at that NT, meriting future examination.

Analyses of population divergence patterns using the VHSV-IVb genome discern similar spatial and temporal patterns as found in our prior partial *G-*gene analyses [75]. These results showed that increased divergence characterized the later time period (2012–2016), as compared to the early (2003–2007) and middle (2008–2011) time periods of VHSV-IVb in the Great Lakes. This pattern is evident in all individual genes as well, suggesting that the *G*-gene depicts overall population trends. During the later time period, VHSV-IVb has occurred more sporadically in smaller, localized outbreaks, which may reflect increased host population immunity and resistance, as predicted by the “Red Queen hypothesis” [49, 50]. Likewise, viral phage Φ2 showed greater genome divergence when exposed to changing hosts versus consistent ones, particularly in genes associated with host immune suppression [103]. On the host side, Alves et al. [104] described genomic changes in immune-related genes of wild rabbit populations following 60 years of exposure to Myoxma Virus (MYXV).

### Evolutionary perspectives for the *Nv*-gene

Our multi-gene phylogenies based on NT and AA sequences reveal little consensus regarding the evolutionary origin of the *Nv* gene. All results support the sister relationship between IHNV and HIRRV. VHSV*-Nv* appears possibly related to the *L*-gene in the NT sequence tree, but pairs with IHNV*-Nv* and SHRV*-Nv* in the AA tree, showing closer relationship to the *G*-gene. SHRV*-Nv* differs the most from other novirhabdoviruses, appearing more closely related to the *G*-gene in the NT tree and to the *P*-gene for the AA tree. This discrepancy might reflect differential functionality, since studies showed that *Nv* is not essential to pathogenicity in SHRV [20, 22]. In contrast, VHSV and IHNV require *Nv* for suppression of the host immune response [19]. We conclude that *Nv* is highly saturated and homoplastic, especially for SHRV.

### Differences in cytopathogenicity

After 24–72 hpi, the 2016 isolates produced less VHSV-IVb RNA and induced higher IFN transcription, compared to the reference 2003 strain (CellC03) (Figure 10). Although viral yields did not significantly differ, by 96 hpi CellC03 produced more active viral particles (Figure 8) and was more cytotoxic at higher MOIs (Figure 7). The genetic changes seen between CellC03 and Cell16a–c may have affected their ability to suppress host response and cause cellular damage. These may contribute to the virus appearing less virulent over time. Cell16b–c (both isolated from E16LB) differ by a single *L-*gene NT and from Cell16a (isolated from ESG16a) by four *L* NTs (one AA). Single nonsynonymous changes can have functional consequences for VHSV. Notably, Chinchilla and Gomez-Casado [26] discerned the function of SNPs in *Nv* by inducing mutations in VHSV-Ia strain FR07-71. Ke et al. [43] found that when four AA *M*-gene mutations were introduced to C03MU, the virus was less able to suppress the transcription of host immune defenses. These data suggest that even subtle changes can lead to altered gene function and viral phenotypic characteristics.

It appears that the oldest VHSV-IVb isolate, C03MU, has remained the most virulent to date. More recent isolates from 2006–16 showed reduced virulence, potentially resulting from virus-host coevolution. Other studies observed similar trends. For example, Imanse et al. [44] examined vcG001 (C03MU) and vcG002 (not included in our analysis), finding faster growth but lower titers in cells exposed to vcG002. Those had similar levels of viral RNA, suggesting that vcG002 was less efficient at producing infective particles. Getchell et al. [45] also found reduced viral load in O06SB, O13GS, and E14GS. In our results, viral RNA production and virulence of Cell16a–c likewise did not significantly differ from CellC03, as evidenced by lower peak virus production in the viral yield assay. Reduced virulence over time has characterized MYXV in Australian rabbits [105], human papillomaviruses [106], and RABV in hyenas [107], leading to a longer infectious period that may aid spread of the virus. Such pattern may similarly characterize VHSV-IVb, meriting further work.

From our findings, we are unable to determine the mechanism underlying the observed reduction in virulence for the 2016 isolates. Reduced virulence might indicate less ability to suppress the host immune response, or it could be that the stronger immune response is the consequence of reduced virulence itself, allowing infected cells more time to produce antiviral factors. Further investigations into the impact of individual SNP changes seen in the 2016 isolates might answer this lingering question. Other investigations observed large impacts on virulence from single nucleotide changes in VHSV-I [27] and Vesicular Stomatitis Virus (VSV) [108]. It is possible that reduced virulence may play a role in aiding transmission.

## Conclusions

As VHSV-IVb nears the end of its second decade of evolution in the Great Lakes region, it has continued to change genetically and in virulence. IVb contains fewer SNPS than have occurred in IVa or I/III, with IVa appearing to evolve the fastest, and rates of IVb and I/III being similar to each other. Our *in vitro* studies show that more recent isolates exhibit reduced virulence, consistent with other findings and field observations. Overall, VHSV-IVb may continue to comprise a threat to fish populations in the Great Lakes region and other waterways, as it continues to evolve. These patterns suggest an overall trend towards persistence at relatively low levels and lesser virulence in host populations over time, in concert with predictions of the Red Queen hypothesis.

## Acknowledgements

Research funding was provided by grants to CAS and DWL, including NSF-DBI-1354806 for “*Gene diversity of the VHS fish virus: Evolution of cellular immune response and pathogenesis*” and USDA-ARS CRIS #3655-31320-002-00D, under specific cooperative agreement #58-3655-9-748, “*VHS fish virus in yellow perch aquaculture*”. We thank S. Edwards for field and laboratory work, N. Marshall for computational and laboratory help, and G. Kurath and J. Winton for providing historical samples. We are grateful to F. Averick, D. Butterfield, F. Calzonetti, K. Czajkowski, A. Elz, T. Fisher, M.E. Hernandez Gonzalez, A. Izzi, M. Krishnamurthy, D. Moorhead, and S. McBride for logistical assistance. This is contribution #4974 from the NOAA Pacific Marine Environmental Laboratory (PMEL).

## Author contributions

CAS conceived the study, and supervised the overall evolutionary study and MDN’s graduate work; CAS and DWL wrote the NSF grant that funded the work; MDN helped fine-tune the concept, collected the data and analyses, designed the figures and tables, and wrote the first draft; DWL and BG oversaw the cell culture and immune response protocols; MDN, CAS, and DWL interpreted the data and wrote the manuscript. All authors revised and edited the manuscript. Final revision, editing, and proof-reading were provided by CAS.

## Data accessibility

All sequences are deposited in NIH GenBank as MK782981–MK783014 (see Table 1). All analyses are deposited in Dryad under XXX.

---

**Supplementary Table A.** Single nucleotide polymorphisms (SNPs) and nonsynonymous changes per individual isolates. *=Group, includes C06NP, C06RB, C06SR, C06YP, C06FD, M08AM, C08LEa, C08LEb, and C09MU.

| Isolate | *N*-gene |  | *P*-gene |  | *M*-gene |  | *G*-gene |  | *Nv*-gene |  | *L*-gene |  | NCDS | Total |  | Average |  | dN/dS |
| --- | --- | --- | --- | --- | --- | --- | --- | --- | --- | --- | --- | --- | --- | --- | --- | --- | --- | --- |
|  | NT | AA | NT | AA | NT | AA | NT | AA | NT | AA | NT | AA |  | NT | AA | NT | AA |  |
| E06FD | 1 | 0 | 0 | 0 | 0 | 0 | 0 | 0 | 0 | 0 | 3 | 0 | 0 | 4 | 0 | 1.6% | 0.0% | 0.000 |
| E06WA | 0 | 0 | 0 | 0 | 1 | 1 | 1 | 1 | 2 | 1 | 2 | 1 | 1 | 7 | 4 | 2.8% | 4.7% | 0.571 |
| E06WBb | 0 | 0 | 0 | 0 | 1 | 1 | 0 | 0 | 0 | 0 | 3 | 1 | 1 | 5 | 2 | 2.0% | 2.4% | 0.400 |
| E06YPa | 0 | 0 | 0 | 0 | 1 | 1 | 2 | 0 | 0 | 0 | 2 | 1 | 1 | 6 | 2 | 2.4% | 2.4 | 0.333 |
| E06SB | 1 | 0 | 0 | 0 | 1 | 1 | 1 | 1 | 0 | 0 | 2 | 1 | 1 | 6 | 3 | 2.4% | 3.5% | 0.500 |
| E06YPb | 0 | 0 | 0 | 0 | 1 | 1 | 0 | 0 | 0 | 0 | 2 | 1 | 0 | 3 | 2 | 1.2% | 2.4% | 0.667 |
| E06YPc | 0 | 0 | 0 | 0 | 1 | 1 | 1 | 1 | 0 | 0 | 2 | 1 | 1 | 5 | 3 | 2.0% | 3.5% | 0.600 |
| E06WBb | 0 | 0 | 1 | 0 | 1 | 1 | 3 | 3 | 0 | 0 | 3 | 1 | 1 | 9 | 5 | 3.6% | 5.9% | 0.556 |
| O06SB | 1 | 0 | 0 | 0 | 2 | 2 | 6 | 3 | 1 | 1 | 11 | 3 | 1 | 22 | 9 | 8.7% | 10.6% | 0.409 |
| C06NP* | 0 | 0 | 0 | 0 | 0 | 0 | 0 | 0 | 0 | 0 | 1 | 1 | 0 | 1 | 1 | 0.4% | 1.2% | 1.000 |
| C06GS | 1 | 0 | 0 | 0 | 2 | 1 | 1 | 1 | 2 | 0 | 3 | 1 | 5 | 14 | 3 | 5.5% | 3.5% | 0.214 |
| B07BG | 0 | 0 | 0 | 0 | 0 | 0 | 2 | 0 | 2 | 1 | 0 | 0 | 3 | 7 | 1 | 2.8% | 1.2% | 0.143 |
| B07PS | 1 | 0 | 0 | 0 | 0 | 0 | 1 | 0 | 0 | 0 | 3 | 2 | 1 | 6 | 2 | 2.4% | 2.4% | 0.333 |
| E07CC | 1 | 0 | 0 | 0 | 1 | 1 | 2 | 1 | 0 | 0 | 3 | 2 | 1 | 8 | 4 | 3.2% | 4.7% | 0.500 |
| E07YPa | 0 | 0 | 0 | 0 | 1 | 1 | 2 | 2 | 1 | 1 | 2 | 1 | 2 | 8 | 5 | 3.2 | 5.9% | 0.625 |
| E07YPb | 0 | 0 | 0 | 0 | 1 | 1 | 2 | 2 | 0 | 0 | 3 | 2 | 2 | 8 | 5 | 3.2 | 5.9 | 0.625 |
| M07SB | 1 | 0 | 0 | 0 | 1 | 1 | 3 | 3 | 0 | 0 | 3 | 1 | 0 | 8 | 5 | 3.2 | 5.9 | 0.625 |
| M08RB | 1 | 1 | 0 | 0 | 0 | 0 | 7 | 4 | 0 | 0 | 6 | 2 | 0 | 14 | 7 | 5.5% | 8.2% | 0.500 |
| E08ES | 2 | 2 | 2 | 2 | 1 | 1 | 0 | 0 | 0 | 0 | 8 | 3 | 1 | 14 | 8 | 5.5% | 9.4% | 0.571 |
| E08FDa | 0 | 0 | 0 | 0 | 1 | 1 | 0 | 0 | 0 | 0 | 3 | 1 | 1 | 5 | 2 | 2.0% | 2.4% | 0.400 |
| E08FDb | 0 | 0 | 0 | 0 | 1 | 1 | 1 | 0 | 0 | 0 | 4 | 1 | 1 | 7 | 2 | 2.8% | 2.4 | 0.286 |
| M08YP | 1 | 1 | 0 | 0 | 1 | 0 | 2 | 1 | 0 | 0 | 8 | 4 | 0 | 12 | 6 | 4.7% | 7.1% | 0.500 |
| M11YP | 0 | 0 | 1 | 1 | 1 | 1 | 0 | 0 | 2 | 0 | 3 | 2 | 2 | 9 | 4 | 3.6% | 4.7% | 0.444 |
| E12FD | 5 | 2 | 2 | 1 | 2 | 1 | 1 | 1 | 1 | 1 | 18 | 6 | 6 | 35 | 12 | 13.8% | 14.1% | 0.343 |
| O13GS | 4 | 1 | 2 | 0 | 2 | 1 | 8 | 3 | 0 | 0 | 9 | 2 | 4 | 29 | 7 | 11.5% | 8.2% | 0.241 |
| E14GS | 2 | 0 | 1 | 0 | 1 | 1 | 6 | 2 | 0 | 0 | 15 | 5 | 2 | 27 | 8 | 10.7% | 9.4% | 0.296 |
| E15RG | 4 | 2 | 2 | 0 | 1 | 0 | 8 | 4 | 1 | 0 | 9 | 3 | 4 | 29 | 9 | 11.5% | 10.6% | 0.310 |
| E16GSa | 3 | 1 | 3 | 0 | 1 | 1 | 7 | 7 | 2 | 1 | 16 | 5 | 6 | 38 | 15 | 15.0% | 17.6% | 0.395 |
| E16GSb | 3 | 1 | 3 | 0 | 1 | 1 | 6 | 6 | 2 | 1 | 16 | 5 | 6 | 37 | 14 | 14.6% | 16.5% | 0.378 |
| E16GSd | 3 | 2 | 3 | 0 | 1 | 1 | 3 | 3 | 2 | 2 | 4 | 1 | 3 | 19 | 9 | 7.5% | 10.6% | 0.474 |
| E16GSd | 3 | 1 | 3 | 0 | 0 | 0 | 3 | 3 | 2 | 2 | 16 | 5 | 5 | 32 | 11 | 12.6% | 12.9% | 0.344 |
| E16GSe | 4 | 2 | 3 | 0 | 0 | 0 | 5 | 5 | 2 | 1 | 16 | 5 | 6 | 36 | 13 | 14.2% | 15.3% | 0.361 |
| M16RGa | 2 | 1 | 1 | 0 | 4 | 1 | 1 | 1 | 1 | 0 | 4 | 1 | 1 | 14 | 4 | 5.5% | 4.7% | 0.286 |
| M16RGb | 3 | 1 | 4 | 0 | 3 | 1 | 1 | 1 | 1 | 0 | 6 | 0 | 0 | 18 | 3 | 7.1% | 3.5% | 0.167 |
| Cell16a | 4 | 1 | 3 | 0 | 1 | 1 | 6 | 6 | 2 | 1 | 16 | 5 | 6 | 38 | 14 | 15.0% | 16.5% | 0.368 |
| Cell16b | 4 | 1 | 3 | 0 | 1 | 1 | 6 | 6 | 2 | 1 | 12 | 4 | 6 | 34 | 13 | 13.4% | 15.3% | 0.382 |
| Cell 16c | 4 | 1 | 3 | 0 | 1 | 1 | 6 | 6 | 2 | 1 | 13 | 4 | 6 | 35 | 13 | 13.8% | 15.3 | 0.382 |
| CellC03 | 0 | 0 | 0 | 0 | 1 | 1 | 3 | 1 | 0 | 0 | 0 | 0 | 0 | 4 | 2 | 1.6% | 2.4% | 0.500 |
| **AVERAGES** | 1.553 | 0.553 | 1.053 | 0.105 | 1.053 | 0.789 | 2.816 | 2.053 | 0.789 | 0.395 | 6.579 | 2.211 | 2.289 | 16.132 | 6.105 | 0.064 | 0.072 | 0.422 |

**Sup Fig A.**
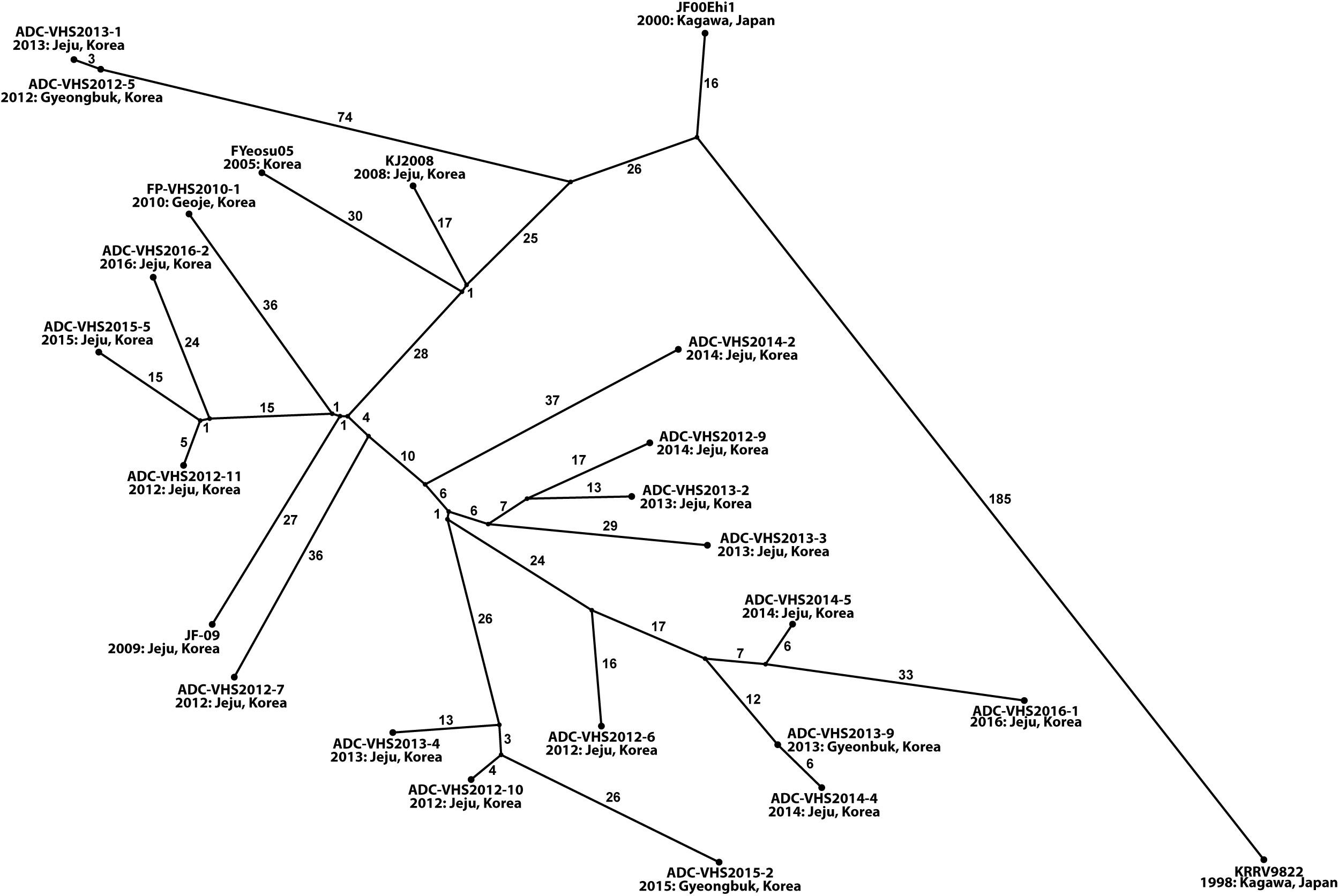
Haplotype network showing genetic relationships among 24 VHSV-IVa full genomes from POPART. Circles sized according to haplotype frequency among isolates. Numbers inside parentheses designate NT differences between each node, unlabeled black circles = hypothesized haplotype steps. Year and location of isolation are below isolate names.

**Sup Fig B.**
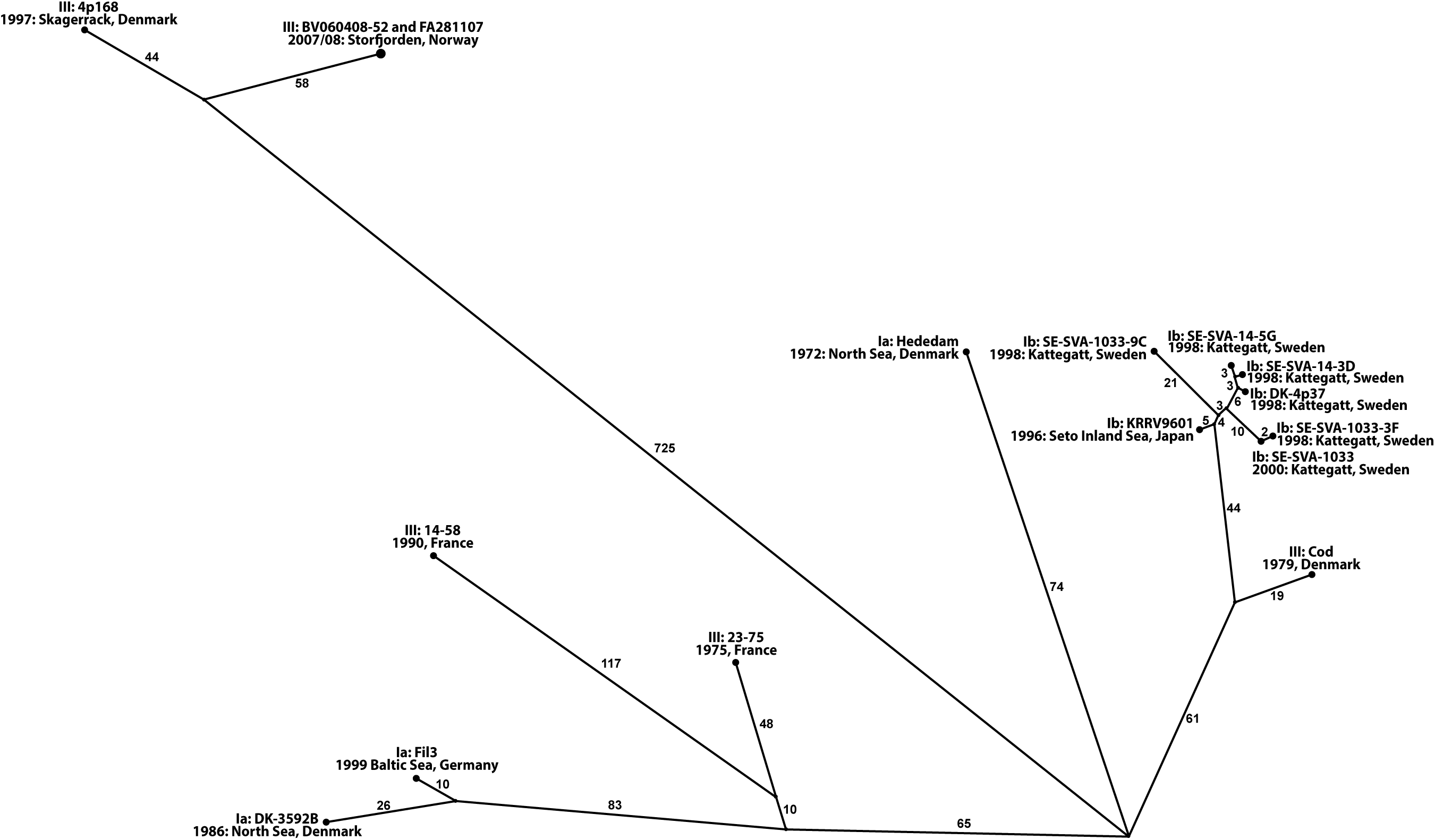
Haplotype network showing genetic relationships among 16 VHSV-Ia, Ib, and III full genomes from POPART. Circles sized according to haplotype frequency among isolates. Numbers inside parentheses designate NT differences between each node. Small, unlabeled black circles = hypothesized haplotype steps. Year and location of isolation are below isolate names.

